# Boiling Acid Mimics Intracellular Giant Virus Genome Release

**DOI:** 10.1101/777854

**Authors:** Jason R. Schrad, Jônatas S. Abrahão, Juliana R. Cortines, Kristin N. Parent

## Abstract

Since their discovery, giant viruses have expanded our understanding of the principles of virology. Due to their gargantuan size and complexity, little is known about the life cycles of these viruses. To answer outstanding questions regarding giant virus infection mechanisms, we set out to determine biomolecular conditions that promote giant virus genome release. We generated four metastable infection intermediates in Samba virus (lineage A *Mimiviridae*) as visualized by cryo-EM, cryo-ET, and SEM. Each of these four intermediates reflects a stage that occurs *in vivo*. We show that these genome release stages are conserved in other, diverse giant viruses. Finally, we identified proteins that are released from Samba and newly discovered Tupanvirus through differential mass spectrometry. Our work revealed the molecular forces that trigger infection are conserved amongst disparate giant viruses. This study is also the first to identify specific proteins released during the initial stages of giant virus infection.

## Introduction

A hallmark of newly discovered giant viruses (GV) is their incredibly complex biology, including gargantuan capsid sizes and large genomes. The sheer size and complexity of these viruses, especially the inclusion of “junk” DNA in the form of introns (Azza et al., 2009; Boratto et al., 2018), challenges the canonical view of viruses as small, streamlined, and efficient killing machines. For example, most GV are larger than 300 nm and many have genomes exceeding 1MB, containing an estimated 1000+ open reading frames (see Table 1 in (Colson et al., 2017)). By contrast, some of the smallest viruses include the porcine circovuris (17 nm capsid, ∼2000 base genome, four proteins, (Dhindwal et al., 2019)) and the human rhinovirus (∼7200 base genome, 30 nm capsid, 11 proteins, (Jacobs et al., 2013)). ∼69% of known viruses encode for less than 10 proteins (Brandes and Linial, 2019), highlighting the complexity of GV and the true extent of our lack of knowledge concerning this new class of viruses.

GV have been isolated from a wide variety of hosts, including amoeba (Aherfi et al., 2016b), animals (Andrade et al., 2018; Andrade et al., 2015; Dornas et al., 2014a; Khan et al., 2007), as well as human and murine cells (Ghigo et al., 2008; Lusi et al., 2017). However, amoebas also infect these creatures, casting doubt on the true viral reservoir. Although GV have been associated with human diseases such as respiratory diseases (Khan et al., 2007; La Scola et al., 2005; Saadi et al., 2013a; Saadi et al., 2013b), inflammatory conditions (Popgeorgiev et al., 2013; Shah et al., 2014), and cancers (Aherfi et al., 2016a), no direct link between GV and human disease has yet been established. Despite an unusually broad host range and pathogenicity, little information is available on how GV access their hosts. Host cell infection usually occurs via phagocytosis (Abrahao et al., 2014; Ghigo et al., 2008). Once phagocytosed, a unique capsid vertex opens which promotes nucleocapsid release and fusion with the phagosomal membrane, ultimately releasing the genome into the host cytoplasm. A pseudo-organelle, called a viral factory, is then formed (Suzan-Monti et al., 2007) and host replication factors are hijacked. The endpoint of GV infections is host cell death and release of new GV progeny into the environment.

GV are ubiquitous (Aherfi et al., 2016b; Andrade et al., 2018) and maintain infectivity in harsh environments such as alkaline lakes (Abrahao et al., 2018), frozen permafrost (Legendre et al., 2014), 3 km deep in the ocean (Abrahao et al., 2018) and dry valleys in Antarctica (Andrade et al., 2018; Kerepesi and Grolmusz, 2017). GV have retained infectivity following exposure to harsh chemicals (Campos et al., 2012), extreme pH and salinity (Abrahao et al., 2018), extreme temperatures (Andrade et al., 2018; Legendre et al., 2014), and are able to persist on hospital equipment (Campos et al., 2012; Dornas et al., 2014b). To survive such extremes, GV have developed incredible capsid stability. Some giant viral capsids can retain infectivity for 30,000 years in permafrost (Legendre et al., 2014; Legendre et al., 2015).

Although capsid stability is beneficial for a virus to persist in harsh environments, it also creates a thermodynamic barrier that must be overcome once a suitable host cell is encountered. Traversing an energy barrier to promote infection and genome transfer into a host cell is not a problem unique to GV; all known viruses must do this to propagate. Strategies and structures used for genome translocation are conserved across viral families. Amongst the tailed dsDNA bacteriophages (*Caudovirales*), tail complexes interact with host receptor proteins to trigger conformational changes in the virion, leading to genome release (Parent et al., 2018). Similarly, many classes of eukaryotic viruses have conserved genome release mechanisms. Most enveloped viruses, including HIV, influenza, Zika virus, and herpesvirus, utilize one of three structurally conserved membrane fusion protein varieties (Harrison, 2015). Non-enveloped viruses, such as rhinovirus, poliovirus, and adenovirus, utilize conserved capsid structures to interact with host receptors to trigger genome uncoating (Suomalainen et al., 2013).

Morphologically, GV virions are either icosahedral, as exemplified by Acanthamoeba polyphaga mimivirus (La Scola et al., 2003), or non-icosahedral typified by Mollivirus and Pithovirus (Legendre et al., 2014; Legendre et al., 2015). Similar to their smaller cousins, GV also share conserved capsid structures that are used during infection. In many GV, the unique capsid vertex provides a gateway for the infection process, but they also provide a mechanism to prevent premature loss of their precious cargo. GV have developed at least two distinct vertex structures to seal the unique vertex until the time is right for infection: “corks” and “starfish”. Non-icosahedral GV tend to utilize one or more cork-like structures to seal their unique capsid locations (Andreani et al., 2016; Legendre et al., 2014; Philippe et al., 2013). These complexes are located flush with the capsid surface. A newly-discovered class of non-icosahedral GV, consisting of members such as Pandoravirus (Legendre et al., 2014) and orpheovirus (Andreani et al., 2017a), contain an ostiole-like structure, distinct from the cork-like structure.

Mimivirus-like icosahedral GV utilize an external proteinaceous seal complex that resembles a five-pointed starfish (Klose et al., 2010; Xiao et al., 2009). These complexes sit at the outermost layer of the capsid at a unique five-fold vertex (called the stargate vertex due to its symmetry and appearance) and prevent it from opening (Xiao et al., 2009). Traditionally, both the unique capsid vertex and the external seal complex have been packaged together and called either the “stargate” or the “starfish”. We will refer to the unique capsid vertex as the stargate and the seal complex as the starfish. Non-mimivirus-like icosahedral GV such as PBCV-1 (Milrot et al., 2017), Faustovirus (Klose et al., 2016), and Pacmanvirus (Andreani et al., 2017b) do not utilize stargate vertices and have evolved alternative genome release strategies.

Starfish structures are found in diverse GV such as mimivirus (Klose et al., 2010; Xiao et al., 2009), Samba virus (SMBV, (Campos et al., 2014; Schrad et al., 2017)), and the newly discovered *Tupanviruses* (Abrahao et al., 2018; Silva et al., 2019), and are more common than the cork-like seals amongst GV. Yet, relatively little is known about the mechanism governing the stargate. The molecular forces and biochemical trigger(s), such as receptor proteins or phagosomal transitions that facilitate stargate opening are unknown. Additionally, the ultimate fate of the starfish remains a mystery; is the complex removed from the capsid *en masse*, or does the complex simply unzip?

The general steps and macroscopic, gross morphological changes that accompany GV infection have been visualized via thin section transmission electron microscopy (TEM) of infected cells (Abrahao et al., 2014; Silva et al., 2019). Following phagocytosis the stargate vertex begins to open between 1-3 hours post infection (Silva et al., 2019), yet, little is known about the specific proteins and biomechanical forces that mediate this process. This knowledge gap is largely due to two factors, the complexity of GV virions and the lack of a robust model system for detailed biochemical and/or biophysical studies. Here, we have created the first *in vitro* model system for studying the choreography that governs GV genome release using SMBV, a member of *Mimiviridae* lineage A (Campos et al., 2014). We were able to trap infection intermediates, identify specific proteins released during the initial stage of stargate opening, and test the efficacy of this technique on other icosahedral GV including a mimivirus variant, M4 (Boyer et al., 2011), Tupanvirus soda lake (TV, (Abrahao et al., 2018)), and Antarctica virus (Andrade et al., 2018). Additionally, our model reveals that members of *Mimiviridae* lineage A unzip their starfish complexes to initiate infection.

## Results and Discussion

### SMBV is Resistant to the Vast Majority of Chemical Treatments

To probe the molecular forces that play a role in SMBV starfish complex stability, we exposed SMBV to treatments known to affect morphology and infectivity in other viruses (Table S1). The effect of each treatment on particle stability was assessed via cryo-EM. Treatments included the denaturants urea (up to 9 M) and guanidinium hydrochloride (up to 6 M), the detergent Triton X-100, organic solvents such as chloroform and DMSO, as well as enzymes including DNase I, bromelain, proteinase K, and lysozyme. None of these treatments resulted in disruption of the SMBV virion, over the baseline of ∼5% spontaneously open SMBV particles as observed under native conditions (Schrad et al., 2017). Two treatments did lead to significantly increased disruption of the stargate vertex: low pH and high temperature (see following sections).

### Electrostatic Interactions are Critical for SMBV Starfish Stability

We hypothesized that pH changes occurring during and after phagocytosis may trigger SMBV stargate opening. Therefore, we dialyzed SMBV particles against different sodium phosphate buffer solutions, ranging in pH from 2-12 (Figure 1A). Particles were visualized via cryo-EM (Figure 2E) and the percent of open particles (POP) was calculated. At and above pH 4, there was no appreciable change in the POP, compared to native (pH 7.4) levels (Figure 2A-D). However, at and below pH 3, ∼60% of the SMBV capsids had opened. While the conditions that produced an increase in SMBV POP (pH ≤ 3) are more acidic than the environment predicted within the amoebal phagosome (Flannagan et al., 2015; German et al., 2013; Lopez and Skaar, 2018), they are similar. Thus, it demonstrates that our *in vitro* results reflect a relevant stage of the GV infection mechanism.

**Figure 1:**
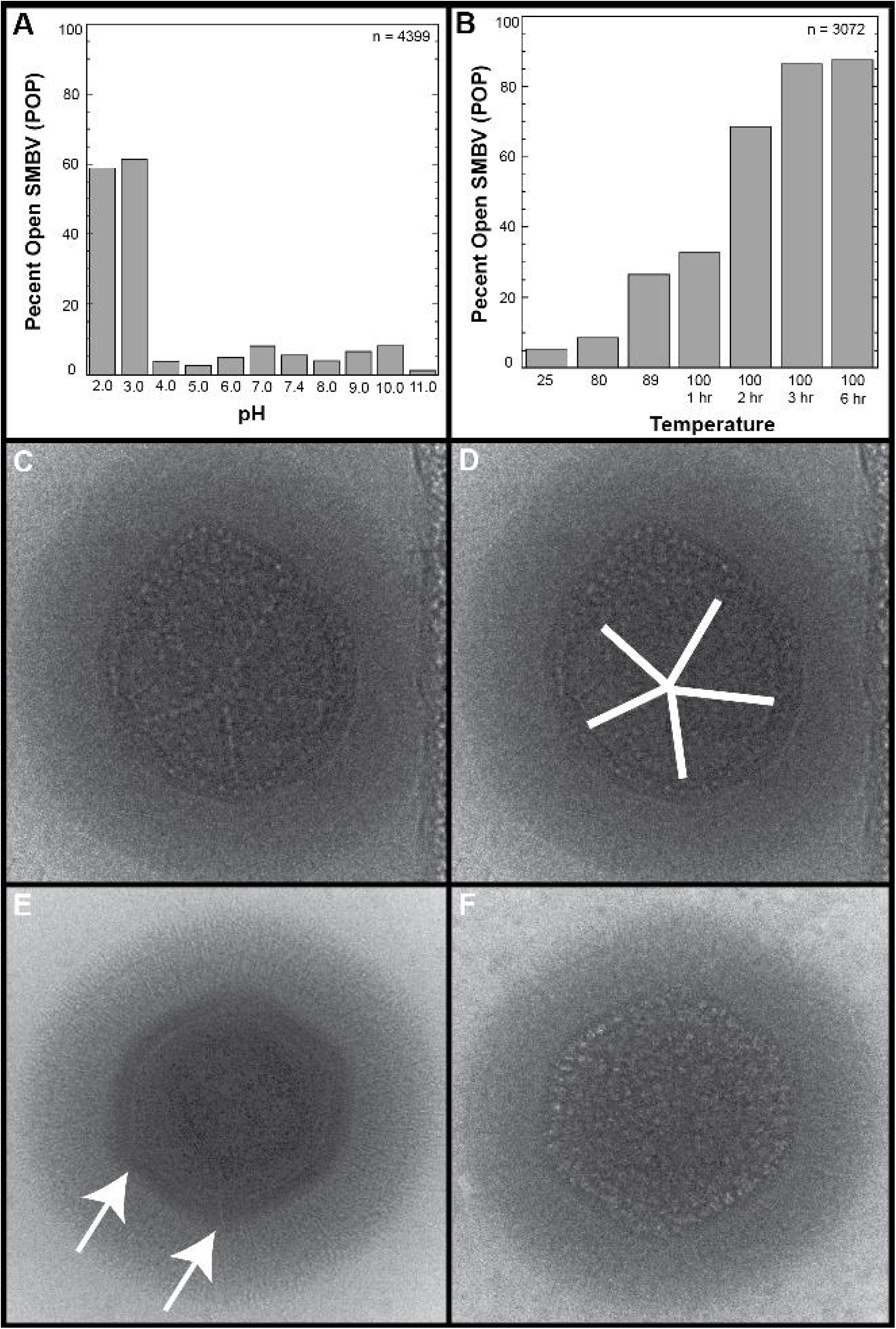
Low pH and High Temperature Triggered an Increase in SMBV POP and Changed the Star-Shaped Radiation Damage Pattern. A) The percentage of open SMBV particles (POP) following treatment at various pH (see Figure S1 and Table S1). B) The POP of SMBV particles incubated at elevated temperatures. C) “Bubblegram” image of a native SMBV particle with a clear star-shaped radiation damage pattern (highlighted in white in D, see Movie S1). E) First exposure in a bubblegram series of a pH 2-treated SMBV particle. The cracked stargate vertex lies in a top-down view. Arrows highlight the slight cracks in the SMBV capsid. F) Final exposure of the bubblegram series begun in E. Note the absence of the star-shaped radiation damage pattern following starfish disruption.

**Figure 2:**
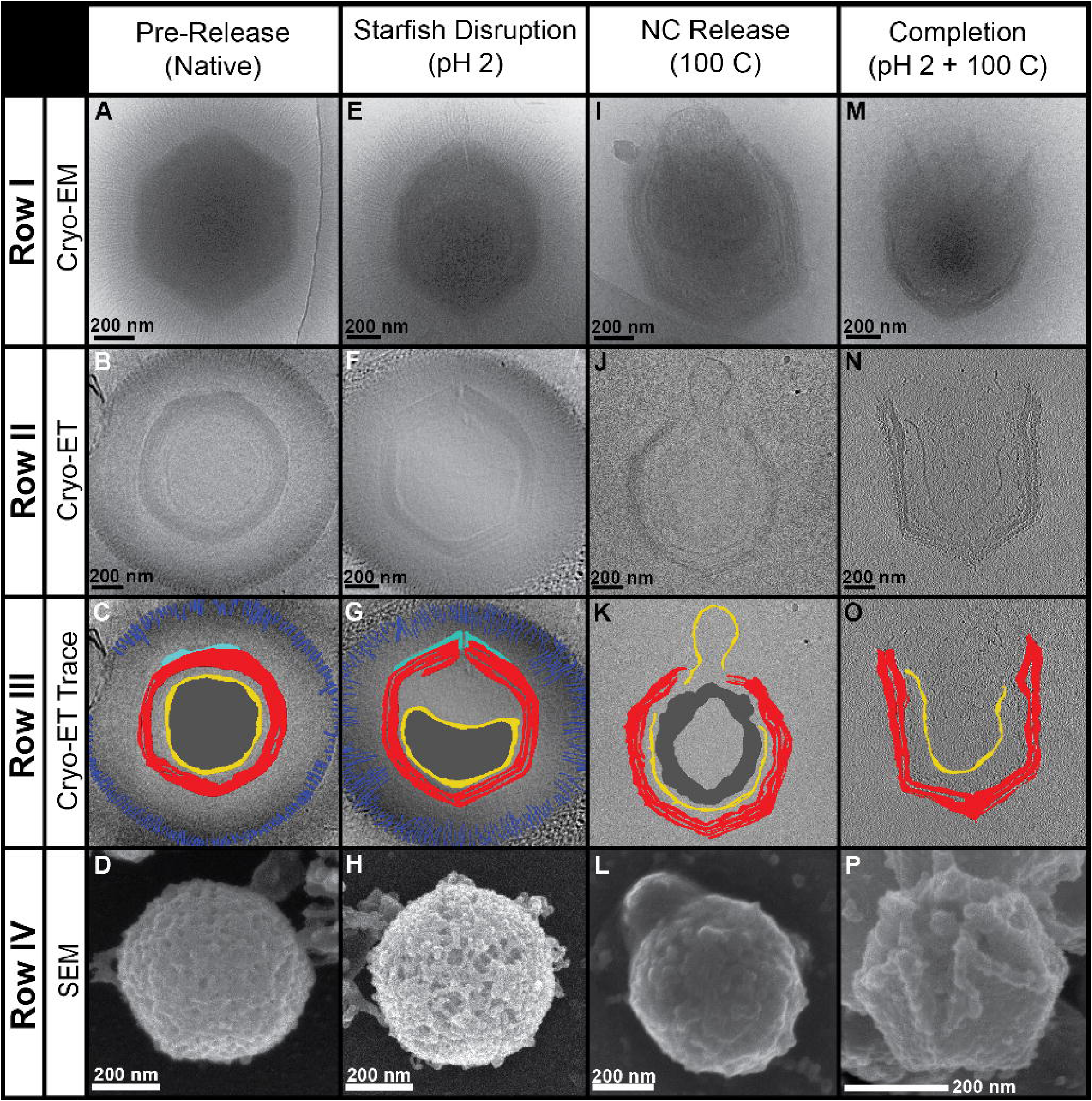
Electron Microscopy of SMBV Genome Release Stages. *Row I)* Two dimensional cryo-electron micrographs of particles following either no treatment (A), or post incubation with pH 2 (E), 100 °C (I), or both pH 2 + 100 °C (M). *Row II)* Central slices (z = 20) of cryo-electron tomograms of particles following either no treatment (B) or post incubation with pH 2 (F) 100 °C (J), or both pH 2 + 100 °C (N). *Row III)* Central slices of cryo-tomograms with key features highlighted. Blue = distal tips of the external fiber layer, Cyan = starfish seal complex, Red = capsid, Yellow = lipid membranes (nucleocapsid), Dark grey = dsDNA. Slices are shown for virions following either no treatment (C) or post incubation with pH 2 (G) 100 °C (K), or both pH 2 + 100 °C (O). *Row IV)* Scanning electron micrographs of particles in various stages of genome release following either no treatment (D) or post incubation with pH 2 (H) 100 °C (L) or both pH 2 + 100 °C (P). See Movies S2-S9 for videos of the tomograms and tilt series. See EMD-20745-20748 for tomogram volumes.

Unlike spontaneously opened GV capsids (Schrad et al., 2017; Xiao et al., 2009; Zauberman et al., 2008), these SMBV capsids were not fully open. Instead, the particles had small, noticeable cracks at one capsid vertex that assumed a star-shaped pattern. The opening of the stargate vertex at low pH is irreversible: SMBV particles returned to neutral pH still displayed star-shaped cracks in their capsids (data not shown). In some particles the extra membrane sac was caught in the process of leaving the capsid through the newly opened vertex (Figure 2E). In other particles, the sac is not visible, suggesting that it had escaped prior to imaging. Release of the sac, also referred to as the viral seed, has been hypothesized in other GV. The viral seed is thought to contain proteins responsible for the formation of the GV viral factory (Mutsafi et al., 2010; Silva et al., 2019; Suzan-Monti et al., 2007). To our knowledge, this is the first study to demonstrate release of the viral seed and to identify some of the proteins that may be released with this complex (below).

We could see that the particles had indeed opened following low pH treatment. Using 2D images alone we could not, however, determine if the starfish complex was released *en masse* or if it remained associated with the capsid. Therefore, we used scanning electron microscopy (SEM) to probe surface features. Unfortunately, SEM images of pH 2-treated SMBV particles (Figure 2H) also did not provide definitive evidence for the presence of the starfish seal as the layer of external fibers blocked access to the capsid surface. We next generated 3D reconstructions of opened SMBV particles through cryo-electron tomography (cryo-ET) (Figure 2F-G, Movie S3, EMD-20747). Tomograms confirm that the stargate vertex, and only the stargate vertex, is open in the pH 2-treated particles. Extra density corresponding to the starfish seal is clearly observed along the edges of the outer capsid layer at the stargate vertex (Movie S3). Therefore, it is likely that at least some, if not all, of the proteins that comprise the starfish seal complex remain attached to the capsid after low pH treatment. The presence of this density in our tomograms suggests that the SMBV starfish likely destabilizes through an “unzipping” mechanism rather than *en masse* release. As low pH treatment is able to trigger stargate vertex opening *in vitro*, we conclude that electrostatic interactions play a very important role in stabilizing this vertex prior to infection.

We turned to “bubblegram” imaging, a cryo-EM imaging technique used for localizing unique features within macromolecular complexes. In this technique, samples are intentionally overexposed to produce beam-induced radiation damage. If there is a unique feature within a complex, hydrogen (H2) gas released as a result of the radiation damaging can become trapped and sometimes produces noticeable “bubbling” in the micrograph. This bubbling can be used to reveal the location and shape of the unique features in viral capsids (Parent et al., 2018) such as bacteriophage ΦKZ inner bodies (Wu et al., 2012) and also ejection proteins in bacteriophage P22 (Wu et al., 2016).

When untreated SMBV particles were exposed to excessive electron radiation many of the particles produced a star-shaped radiation damage pattern (Figure 1C-D, Movie S1). By contrast, pH 2-treated SMBV particles, displayed no star-shaped pattern (Figure 1E-F). As expected, the lack of a star-shaped radiation damage pattern is consistent with the hypothesis that the H2 gas is no longer being trapped in the SMBV virion as the low pH treatment disrupted the stargate vertex seal.

### Increased Entropy is Required for Nucleocapsid Release

Lowering pH alone was insufficient to fully open SMBV particles, indicating that electrostatic interactions are not solely responsible for sealing the stargate. Therefore, we analyzed the effect of temperature on the stability of SMBV particles. We incubated the virions one hour at up to 100 °C, assayed the virions for morphological changes using cryo-EM, and then compared these data to images of particles that had been incubated at room temperature (25 °C). After 1 hour at 100 °C, the POP was ∼33 % (Figure 1B). Following an additional incubation for up to five hours, the POP increased to a maximum of ∼88%.

Unlike low pH, which simply cracks the stargate vertex, higher temperatures resulted in open stargate vertices with nucleocapsids in the process of exiting the virion (Figure 2I-L, Movie S4-S5, EMD-20748). Within these nucleocapsids the DNA appears to have reorganized leaving pockets of seemingly empty space (discussed in greater detail below.) Additionally, much of the external fiber layer is removed (Figure 2I-L, Figure S1) and the extra membrane sac is fully released from these particles. The use of high temperatures could be an alternative GV defibering method to that proposed in (Kuznetsov et al., 2010), especially as this previously described technique did not defiber SMBV particles (data not shown). High temperature induces a conformational change that closely mimics a stage of mimivirus infection seen *in vivo* (see Figure 2-III in (Abrahao et al., 2014)), where the nucleocapsid leaves the capsid and prepares to fuse with the amoebal phagosome membrane. As increased thermal energy induces stargate opening *in vitro*, we conclude that entropic barriers must be overcome during GV stargate opening *in vivo*. In the amoeba, these entropic barriers are likely lowered by interaction with a cellular receptor, although the identity of these receptors is currently not known for any GV.

Following both low pH and high temperature treatment (individually) there were pockets within the SMBV nucleocapsids that appear to be devoid of DNA (Figure 2J-K). These seemingly empty pockets are not visible in the untreated SMBV particles (Figure 2B-C). While it is possible that the void inside of SMBV nucleocapsids could be due to the extreme conditions used, it is more likely that this is biologically relevant. These pockets are only observed in SMBV particles that have begun releasing their genome, suggesting that the DNA may undergo reorganization during this process. The SMBV genome contains various chromosome condensation and histone-like proteins that could be used for this function. Mass spectrometry experiments (described below, and shown in Table 1) suggest that many of these proteins remain with the nucleocapsid after the initial opening stage. Genome reorganization is an important stage of many virus infection processes, including HIV (Freed, 2015) and Adenovirus (Mangel and San Martin, 2014). We hypothesize that genome rearrangement is also important for facilitating GV genome release into the host.

### Disrupting Electrostatic Interactions and Increasing Entropy Results in Complete SMBV Genome Release

Individually, low pH and high temperature had different physical effects on SMBV. These disparate treatments are affecting two different types of biomolecular interactions (electrostatic interactions and entropy, respectively) and each appears to contribute to SMBV virion stability. Therefore, we hypothesized that combining low pH and high temperature might have a compound effect on stargate opening. Again, following treatment the SMBV particle morphology was analyzed via cryo-EM (Figure 2M), cryo-ET (Figure 2N-O), and SEM (Figure 2P). These particles have completed the entire genome release process, as seen by the absence of the nucleocapsid. Additionally, SMBV particles were completely defibered and the internal capsid layer(s) appeared to be less rigid than the outer capsid layer (Figure 2O, Movie S6-9, EMD-20745 & EMD-20746). Once disrupted, the capsid is more electron transparent and apparent connections between the two capsid layers were now visible in the tomograms (Figure 2N, Movies S7 & S9). Anchoring/tethering proteins that connect these two capsid layers may play a role in the extraordinary capsid stability of GV.

SEM of dual treated SMBV particles (Figure 2P) provides further evidence for the fate of the starfish seal. Particles treated with both low pH and high temperature clearly contain extra density around the edges of the stargate vertex, corresponding to the starfish seal. This extra density is consistent with our cryo-ET data described above where rather than completely dissociating from the capsid *en masse*, the starfish seal unzips to allow the stargate to open while still retaining contacts with the capsid.

### Molecular Forces That Stabilize the SMBV Stargate Vertex are Conserved Amongst Diverse GV

We tested the effects of a combination of pH and temperature on three other GV (from two distinct *Mimiviridae* lineages); Antarctica virus ((Andrade et al., 2018), *Mimivirus A*), TV (Abrahao et al., 2018), and mimivirus M4 ((Boyer et al., 2011), *Mimivirus A*)). Following treatment, each virus was characterized via cryo-EM (data not shown) and SEM (Figure 3). Similar to SMBV, all three GV had opened their stargate vertices and released their nucleocapsids after being boiled in acid. All three GV also appeared to lose the majority of their fibers during treatment. All four of the GV tested in this study had fully open stargate vertices following low pH and high temperature treatment. While all four viruses analyzed here are mimivirus-like icosahedral GV, these viruses encompass two separate GV clades belonging to the *Mimiviridae* family: of the genus *Mimivirus* (SMBV, M4, Antarctica) and the proposed genus *Tupanvirus* (TV, (Rodrigues et al., 2019b)). These data strongly indicate that the general forces that stabilize virions and facilitate infection are conserved among distantly related amoeba-infecting members of *Mimiviridae*.

**Figure 3:**
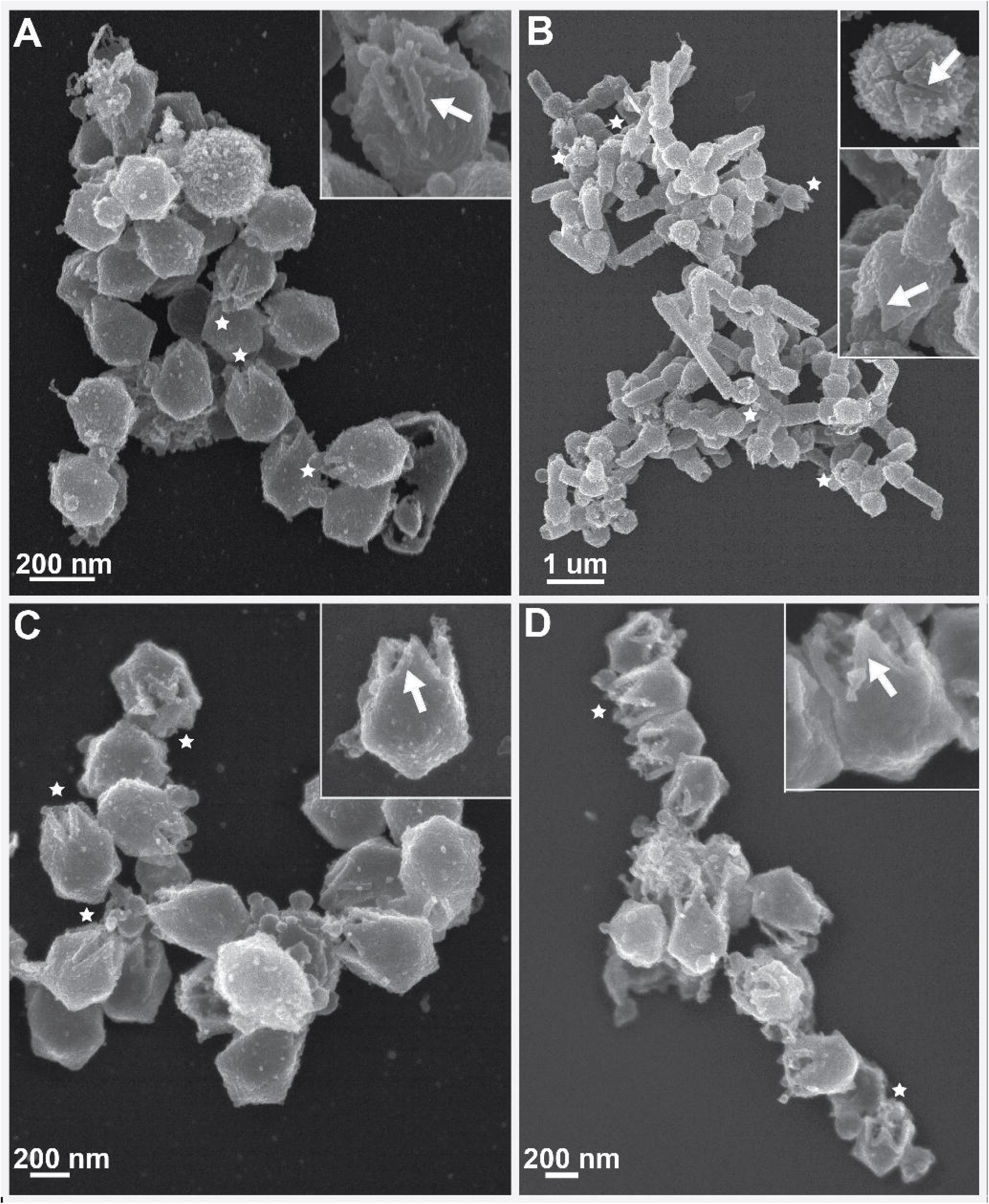
Post Genome Release Particles From Four GV. Scanning electron micrographs of low pH and high temperature-treated A) SMBV, B) TV, C) Antarctica virus, and D) mimivirus particles. Inserts demonstrate enlarged views highlighting capsids where either clear retention of the starfish seal can be seen in SMBV, mimivirus, and Antarctica particles or the lack of starfish seal retention can be seen in TV. Asterisks in the main panels depict selected particles with clearly visible open stargate vertices.

Although the general forces appear to be highly conserved, some specific mechanisms of starfish disruption are likely conserved only within distinct lineages. In our SEM data, Antarctica and mimivirus particles (Figure 3A & 3D, respectively) displayed density along the edges of the open stargate vertices, similar to the density seen in SMBV (Figure 2P, 3C). The presence of this extra density suggests that, like SMBV, the Antarctica and mimivirus starfish complexes unzip to facilitate stargate opening and genome release. TV, on the other hand, does not display this extra density (Figure 3B), suggesting that the TV starfish may completely dissociate from the capsid *en masse* during infection. TV particles also appear to fully open their stargate vertices following low pH treatment alone (data not shown). In total, our data suggest that the mechanism of seal complex unzipping may be conserved amongst *Mimiviridae* with slight deviations present between the *Mimiviruses* and the proposed *Tupanvirus* genus.

GV have changed our canonical view of virology, defying the previously known limits of capsid sizes and stabilities. Giantism is known to cause developmental and structural problems for higher organisms, such as humans (Nabarro, 1987), but GV have evolved a common stargate vertex and accompanying stabilization mechanisms to counteract these issues. The description of a new GV genome release strategy signifies another paradigm shift in our understanding of virology. As mentioned previously, smaller viruses tend to share conserved genome release mechanisms. This conservation can be observed within viral families such as *Flaviviridae* (fusion proteins (Apellaniz et al., 2014)), *Caudovirales* (tail complexes (Parent et al., 2018)), or *Orthomyxoviridae* and *Paramyxoviridae* glycoproteins (Kordyukova, 2017). This conservation also occurs across viral kingdoms. The Herpesvirus portal complex shares structural similarity with many bacteriophage portal proteins (McElwee et al., 2018; Newcomb et al., 2001) and the Adenovirus spike protein is homologous with the bacteriophage Sf6 tail needle knob protein (Parent et al., 2012). GV have eschewed all of these known genome release structures and appear to have forged their own mechanisms, as exemplified by the common stargate mechanism.

### Numerous Proteins are Released From GV Capsids During Stargate Opening

As obvious morphological changes occurred in the GV capsids during low pH and high temperature treatments, we hypothesized that proteins were likely released from the capsids at each of these stages. We analyzed proteins that remained within the SMBV and TV capsids and proteins liberated from the capsids after each treatment. We used four conditions, native virions (pH 7.4, room temperature), low pH (pH 2, room temperature), high temperature (pH 7, 100 °C), and combined (pH 2, 100 °C). We then performed pellet/supernatant separations to physically separate the virions and released proteins. Following separation, we analyzed the contents of each sample via SDS-PAGE (Figure 4). A sample preparation scheme for these experiments can be seen in Figure S2. Antarctica virus and mimivirus both showed a similar banding pattern as SMBV (data not shown). We did not perform MS experiments with these viruses as there is no annotated Antarctica virus genome and mimivirus and SMBV are highly similar (Campos et al., 2012).

**Figure 4:**
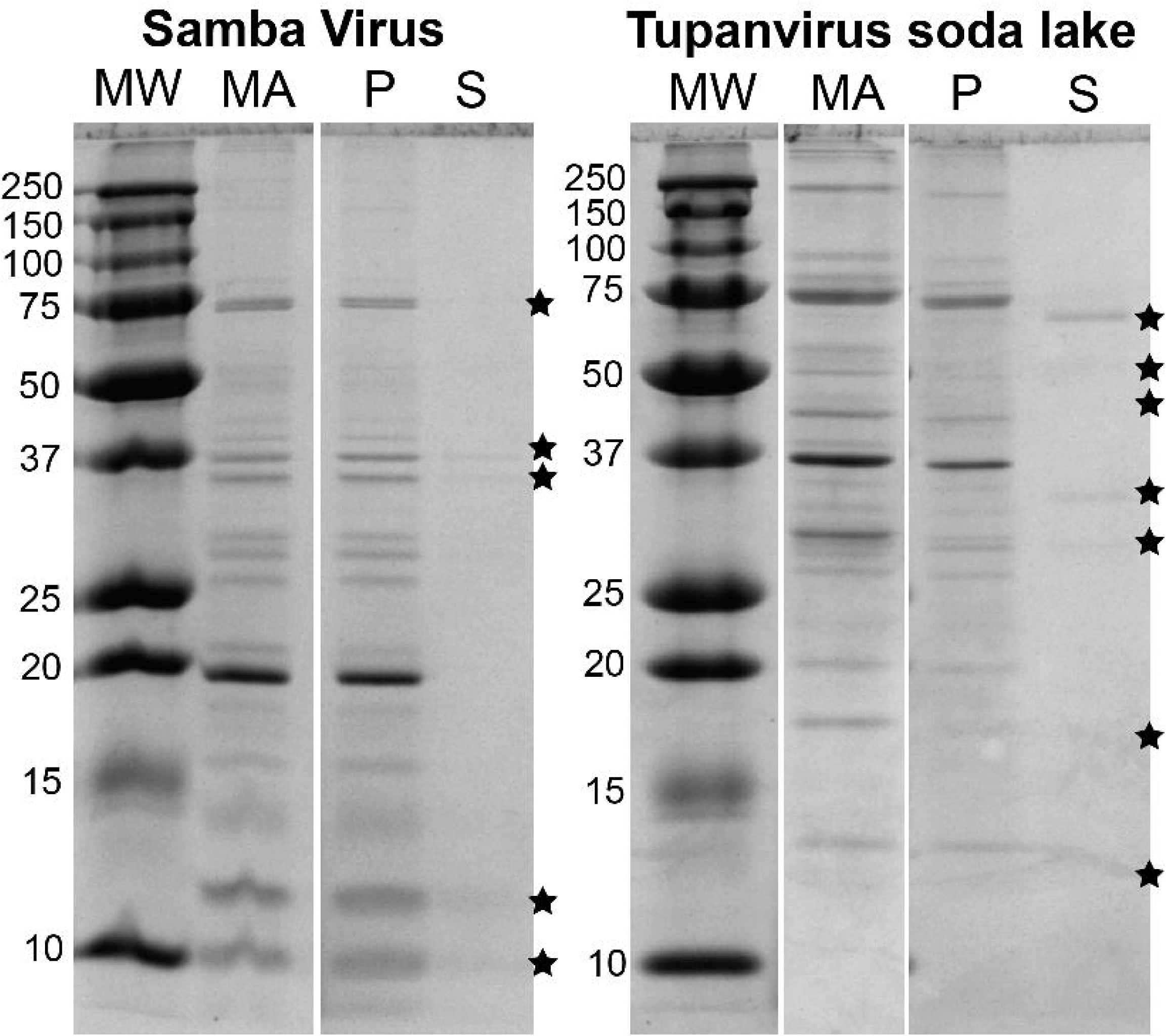
SDS-PAGE of pH 2-Treated SMBV and TV. SDS-PAGE Bands of SMBV and TV. MA = <u>M</u>aterial <u>A</u>pplied (untreated viral particles), P = pellets from pH 2-treated virions, S = supernatants from pH 2-treated virions. Visible bands of proteins released into the supernatant are highlighted with asterisks. See Figure S2 for the sample preparation scheme.

For both SMBV and TV, distinct proteins were released from the capsid following low pH treatment. Some of these proteins align with proteins in the native capsid (pellet) lane, suggesting they had been released from the capsid without significant modification/cleavage. Other proteins, especially in the TV sample, did not match proteins in the native capsid lane. These bands likely represent proteins that were cleaved during low pH treatment. For both viruses, the native supernatant lanes did not contain any visible protein bands. When the particles were incubated at 100 °C (with or without prior pH 2 treatment) it appeared that the majority of proteins were proteolytically cleaved and appeared as a continuous smear on the gel (data not shown) preventing detailed analysis of these samples.

### Identifying the Proteins Released From SMBV and TV Virions at the Initiation of Infection

To characterize the proteins released during the initial stages of GV infection, we used mass spectrometry (MS). Initially, we focused on in-gel digestion of bands from the pH 2-treated SMBV and TV supernatant samples. The low pH-treated particles mimic the beginning of the GV infection process, as the stargate vertex begins to open and the extra membrane sac leaves the capsid. Trypsinized fragments were analyzed via LC/MS/MS and the resultant peptides were compared to published SMBV and TV genome sequences (GenBank KF959826.2 & KY523104.1, respectively) as well as the *A. castellanii* genome (GenBank KB007974.1) to identify any contaminating host proteins from our analysis. The *A. castellanii* actin protein was retained within these results, as this protein is known to play a role in the infection and genome release processes of Iridoviruses (Huang et al., 2018). From this initial experiment, we identified 48 SMBV and 26 TV proteins that are released from the virion following low pH treatment. These proteins are labeled with a (+) in the “Band” column of Table 1.

Excising visible gel bands for MS analysis has the potential to miss proteins within the sample: some bands may be too faint to detect, some proteins may be too large or too small to be fully resolved or extracted, etc. Therefore, we also analyzed SMBV and TV samples using shotgun proteomics to maximize coverage in our study. We analyzed low pH pellet and supernatant samples, as well as the untreated virus using the sample preparation scheme shown in Figure S2. From this experiment we identified 43 SMBV proteins and 37 TV proteins ((+) in the “Shotgun” column of Table 1 and Table S3). Of these proteins, 5 SMBV proteins and 7 TV proteins were previously identified from analysis of the gel bands.

In total, 86 SMBV proteins and 56 TV proteins were identified as having been released from the capsids at low pH. TV was isolated from an environment with high salinity and alkaline pH (9-12, (Abrahao et al., 2018)). SMBV, on the other hand, was isolated from a tributary of the Amazon River, a relatively neutral environment. Due to its location, TV had to evolve pH stability into its capsid to a greater extent than SMBV. While TV was originally isolated from a basic environment some of the strategies that the virus could have developed to stabilize its proteins, such as using a higher percentage of non-polar amino acids, could also stabilize the proteins at low pH.

187 and 169 total proteins were identified within the untreated mature virions of TV and SMBV, respectively (Figure 5). To identify proteins of interest (those that had been released), we calculated the percent of the total peptide signal for each protein. We compared these percentages across the three samples, specifically looking at the ratios of supe:MA (<u>M</u>aterial <u>A</u>pplied) and pellet:MA. Proteins where the supe:MA > 1 were enriched in the treated supernatant sample, indicating that they had been released from the capsids. These proteins are identified with a (+) in the “Up in Supe” column of Table 1. Conversely, proteins with pellet:MA < 1 were less abundant in the treated pellet than the native particles, and likely also released. These proteins are identified with a (+) in the “Down in Pellet” column of Table 1. Proteins that are enriched in the supernatant samples are definitely released from the GV capsids, as no proteins were identified in the untreated supernatant samples (data not shown). Proteins that are depleted in the pellet samples are also likely released from the GV particles, although it is unlikely that any of these proteins are completely absent from the pellet samples (see POP in Figure 1A).

**Figure 5:**
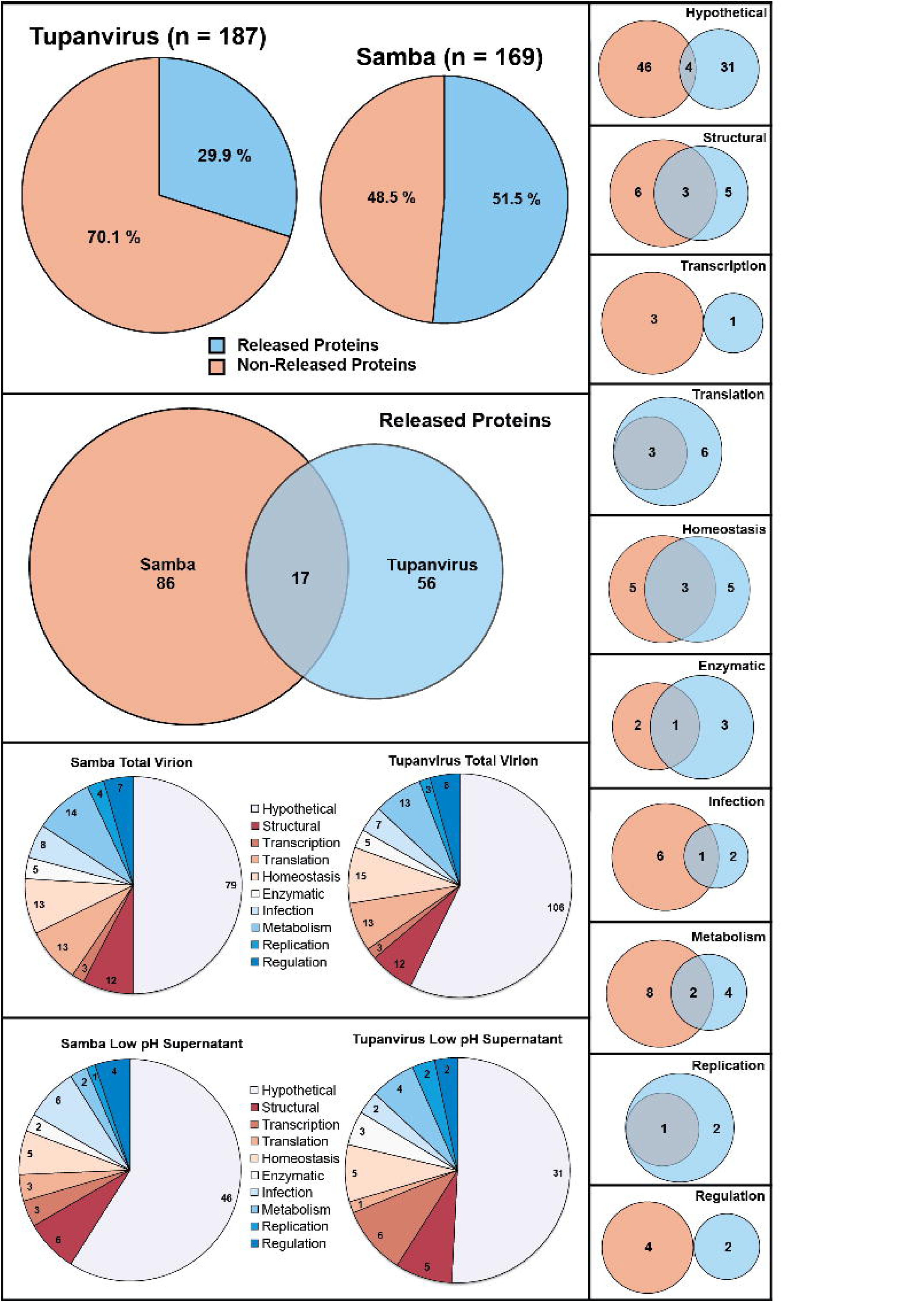
Comparison of Proteins Released by SMBV and TV Soda Lake. Venn diagrams comparing the total protein content (A-B) and proteins released following low pH treatment (C) of SMBV (Red) and TV (Blue) particles. The homology present within these protein sets is depicted in panels D-N. See Tables S2 for hypothetical proteins with predicted transmembrane domains and Table S3 for the relative abundance of individual proteins in each the untreated particles and the treated pellet and supernatant samples.

SMBV releases a higher number and percentage of these proteins (86, 51.5%) than TV particles (56, 29.9%). Putative functions for the released proteins were determined via 1) previous annotation (Abrahao et al., 2018; Campos et al., 2014), 2) NCBI BLAST analysis, 3) HHBLITS analysis (Remmert et al., 2011), 4) InterPro functional prediction (Mitchell et al., 2019), and 5) PSIPRED domain prediction using the DomPred functionality (Buchan and Jones, 2019; Jones and Cozzetto, 2015). Released proteins for each virus were separated into the following 10 categories: Hypothetical, Structural, Transcription, Translation, Homeostasis, Enzymatic, Infection, Metabolism, Replication, and Regulation (Figure 5B-N).

The majority of the proteins released for each virus (53% for SMBV, 55% for TV) are hypothetical proteins or proteins with unknown function. 17 of the proteins released by the two viruses displayed obvious homology between SMBV and TV (BLAST results or functional homology prediction). All of the released SMBV proteins predicted to be involved in both Translation and Replication had homologues amongst the released TV proteins. The proteins predicted to be involved in Transcription and Regulation, on the other hand, did not show any readily apparent homology. The homology between the released TV and SMBV proteins in general and within each category can be found in Figure 5.

### Expected Protein Types are Released from SMBV and TV Virions During Genome Release

GV need to carry out the same basic stages of the viral life cycle as their smaller cousins to replicate. Common stages include genome translocation into the host cell, blocking host replication, hijacking host machinery to make viral proteins, and making new viral proteins (Baker et al., 1999). Both SMBV and TV likely release proteins that are predicted to perform these functions, as many smaller viruses release whole proteins or peptides to facilitate this function (Manning et al., 2018; Parent et al., 2018; Wu et al., 2016). Hypothetical or unknown function proteins released from GV particles likely aid in performing these critical functions as many of them are released during the initial phase of opening. Aside from identifying the putative functions for the hypothetical proteins discussed below, determining the specific function of these proteins lies beyond the scope of this study.

Before the virus is able to hijack the host machinery and begin replication, it must enter the host cell and translocate its genome across the phagosomal membrane into the cytoplasm. SMBV releases putative membrane proteins, such as a virion associated membrane protein (AHJ40731.1) and as well as hypothetical proteins with predicted transmembrane domains that may play a role in membrane fusion (“H/TM” in Table 1). Therefore, the results of this study help to assign putative roles to many proteins with previously unknown function, highlighting the power of this new method.

Additionally, both SMBV and TV release proteins predicted to play a role in a Ubiquitin-Proteasome degradation pathway (UPP, delineated by ^c^ in Table 1). These proteins are known to facilitate genome release in other viruses including the large, but not quite giant Iridoviruses (Huang et al., 2018) and Herpesviruses (Greene et al., 2012). In Iridovirus infection, the UPP is coupled with metabolic, cytoskeletal, macromolecule biosynthesis, and signal transduction proteins to facilitate infection (Huang et al., 2018). Proteins predicted to carry out these functions are released from both the SMBV and TV virions alongside the UPP-related proteins (^e^ in Table 1).

Following genome translocation, the virus forces the cell machinery to transition from making new cellular products to making viral components. Both SMBV and TV release various subunits of a DNA-dependent RNA polymerase (SMBV: AHJ39967.2, AHJ40151.2, AHJ40172.1; TV: AUL78016.1, AUL78362.1, AUL78368.1, AUL78302.1). This series of proteins is critical for the lifecycle of the virus as it directs the cellular machinery of the host to recognize viral DNA *in lieu* of cellular DNA. These proteins, especially the various DNA-dependent RNA polymerases, may play a role in transcription as hypothesized to occur following stargate opening but before nucleocapsid release (Mutsafi et al., 2014) Additional proteins in this category likely include some of the metabolic proteins released by the viruses, especially the catabolic proteins that may play a role in degrading host defenses and machinery. These proteins include a SMBV thiol protease (AMK61869.1), a SMBV amine oxidase (AHJ39955.1), and a hypothetical TV protein with a predicted inosine/uridine-favoring nucleoside hydrolase domain (AUL71835.1). Aside from these RNA polymerase subunits, both TV and SMBV release proteins that facilitate transcription. SMBV releases a poly (A) polymerase (AHJ40056.1), an mRNA-capping enzyme (AHJ40083.1), and an anaerobic transcription regulator (AMK61903.1). TV releases an SNF2 family helicase (AUL77941.1), an ATP-dependent RNA helicase (AUL77829.1), and a mimivirus-like elongation factor (AUL78714.1).

Many of the proteins we identified matched proteins that one would expect to be released during the initial stages of viral infection and greatly supports our hypothesis that the *in vitro* stages generated in this study are reflective of those that occur *in vivo*. These data provide new insights into GV biology and ultimately lead to our proposed model (see next sections).

### SMBV and TV Also Release Novel Proteins During Stargate Opening

SMBV and TV also release proteins that are relatively uncommon amongst viruses. These proteins include metal-binding homeostasis proteins as well as chemotaxis-regulating proteins.

Our mass spectrometry data conclusively show that both SMBV and TV release proteins that are predicted to play a role in maintaining homeostasis (Figure 5E). Many of these proteins are predicted to have redox activity, protecting the virus and its cargo from reactive oxygen species (ROS) that can be found in the host phagosome (Flannagan et al., 2015). These proteins include several thioredoxin-like or thioredoxin domain-containing proteins (<u>SMBV:</u> AHJ40071.1, AHJ40129.2; <u>TV:</u> AUL77963.1) and glutaredoxins (<u>SMBV:</u> AMK618100.1; <u>TV:</u> AUL78724.1). TV releases a catalase protein (AUL78097.1) as well a glyoxylase (AUL78134.1) while SMBV releases a prolyl 4-hydroxylase (AMK61959.1). These proteins are also projected to protect the GV from ROS during the infection process. Here we show that these proteins are indeed released very early in the infection process.

Redox-active proteins are also thought to play an important role in protecting the viruses from the harsh conditions present in the host phagosome. During phagocytosis amoebal phagosomes drop to ∼pH 4 (not low enough to trigger stargate opening), but they are also inundated with metals (like Cu and Zn) and reactive oxygen species (German et al., 2013; Lopez and Skaar, 2018). Both viruses release metal-binding proteins (identified by ^d^ in Table 1) including SMBV’s lanosterol demethylase (AHJ40393.1) --a cytochrome p450-like protein-- and prolyl 4-hydroxylase (AMK61959.1) and TV’s mg709 (AUL77661.1) --a putative prolyl 4-hydroxylase with iron ion binding capabilities-- and Cu-Zn superoxide dismutase (AUL78503.1). It is likely that these proteins, in conjunction with the ROS-mitigating proteins described above, allow these viruses to survive the onslaught of low pH, high ROS, and high metal concentration found inside of the host phagosomes. We also note that the low pH of the phagosomes is similar to the low pH used in our in vitro assay, likely reflecting a physiologically relevant stage that describes GV infection mechanisms.

While *Tupanvirus* infection is hypothesized to occur through phagocytosis (Silva et al., 2019), no biological data has yet been provided to substantiate said hypothesis. This proposal stems from visualization of phagocytosis of TV by *Vermamoeba vermiformis* and subsequent TV stargate opening via thin-section TEM. Thin section TEM, embedding biological samples within epoxy resin then slicing thin, electron translucent sections off of the block, is prone to structural artifacts (Baker et al., 1999). Therefore, it is critical that any hypotheses generated from thin section TEM imaging are supported by data from another technique. The release of proteins capable of mitigating the harsh environment of the amoebal phagosome provides biological evidence to support this hypothesis.

SMBV and TV also contain proteins that are predicted to regulate chemotaxis. SMBV releases a chemotaxis protein (AHJ40337.1) that shares homology with the putative chemotaxis protein CheD found in mimivirus (AKI80461.1) and TV (AUL78687.1). CheD proteins regulate chemotaxis via deamidation of chemotaxis receptors (InterPro). TV has been shown to shut down host chemotaxis (Oliveira et al., 2019; Rodrigues et al., 2019a) and it is likely that these CheD-like chemotaxis regulation proteins are involved in this process. While TV does contain a CheD-like chemotaxis protein that was identified in the total virion MS data, this protein was not present following low pH treatment.

### Making Some Sense of the Myriad Hypothetical Proteins in the SMBV and TV Proteome

Of the 356 proteins identified in the total virion MS for both SMBV and TV, ∼52% (Figure 5B) were annotated as being hypothetical proteins, low complexity proteins (SMBV), or ORFans (an open reading frame that is not found in other reported genomes). In SMBV, 77 of these proteins were released (46 proteins) and in TV 31 proteins are released following low pH treatment. As these proteins are released from the GV virion during the initial stages of the genome release process, we hypothesize that these proteins play a role in either the infection process (phagosome survival, membrane fusion, etc.) or in the beginning stages of replication. Hypothetical proteins with additional functional information predicted via BLAST, HHBLIST, PSIPRED, or InterPro are listed in Table S2. Interestingly, only four of the hypothetical proteins released by the two GV share homology when analyzed via BLAST and HHBLITS, suggesting that while related, SMBV and TV have significant evolutionary divergence.

### Opening the Stargate to New Avenues of GV Research

By modulating temperature and pH we were able to mimic four unique, and metastable stages of the GV genome release process (Figure 6). GV particles that mimic these genome release stages have been seen in previous experiments (Abrahao et al., 2018; Schrad et al., 2017; Xiao et al., 2009; Zauberman et al., 2008), although previously visualization of these particles relied on finding the “one-in-a-million” particle in the correct state. We are now able to mimic GV genome release stages reliably and with high frequency. Additionally, these conditions forgo the need to synchronize infection and trap GV particles in phagosomes at very specific times to generate the condition of interest. Eschewing the host cell may limit specific avenues of study, such as searching for a host receptor(s), but it dramatically simplifies any studies aimed at the virus and the changes it undergoes during the genome release process.

**Figure 6:**
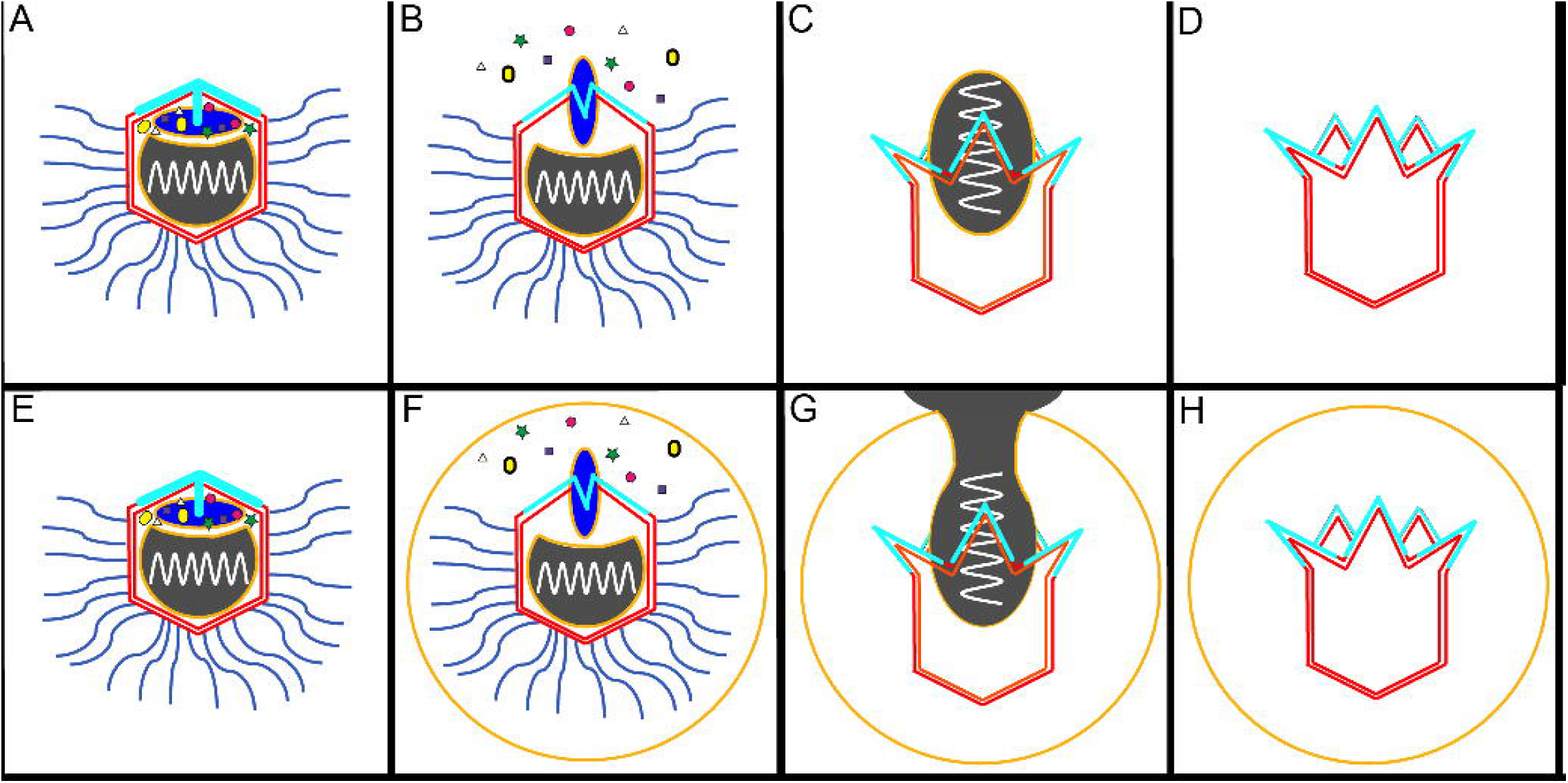
Cartoon Model of Giant Virus Genome Release Stages. Schematic of at least four distinct stages of the GV genome release process as identified in this study. A) <u>Native, intact virions</u>. B) <u>Disruption of the Starfish Seal (Initiation):</u> Particles with stargate vertices that are beginning to open. C) <u>Nucleocapsid Release:</u> Particles with fully open stargate vertices that are in the process of releasing the nucleocapsid from the capsid. D) <u>Fully Released (Completion):</u> Particles that have completed the genome release process. Coloration matches scheme used in Figure 2. The top row depicts particle states as induced *in vitro.* The bottom row corresponds to analogous structures seen in thin section micrographs of infected cells (Aherfi et al., 2016b). The orange circles in panels F, G, and H correspond to the phagosomal membrane.

**Table 1:**
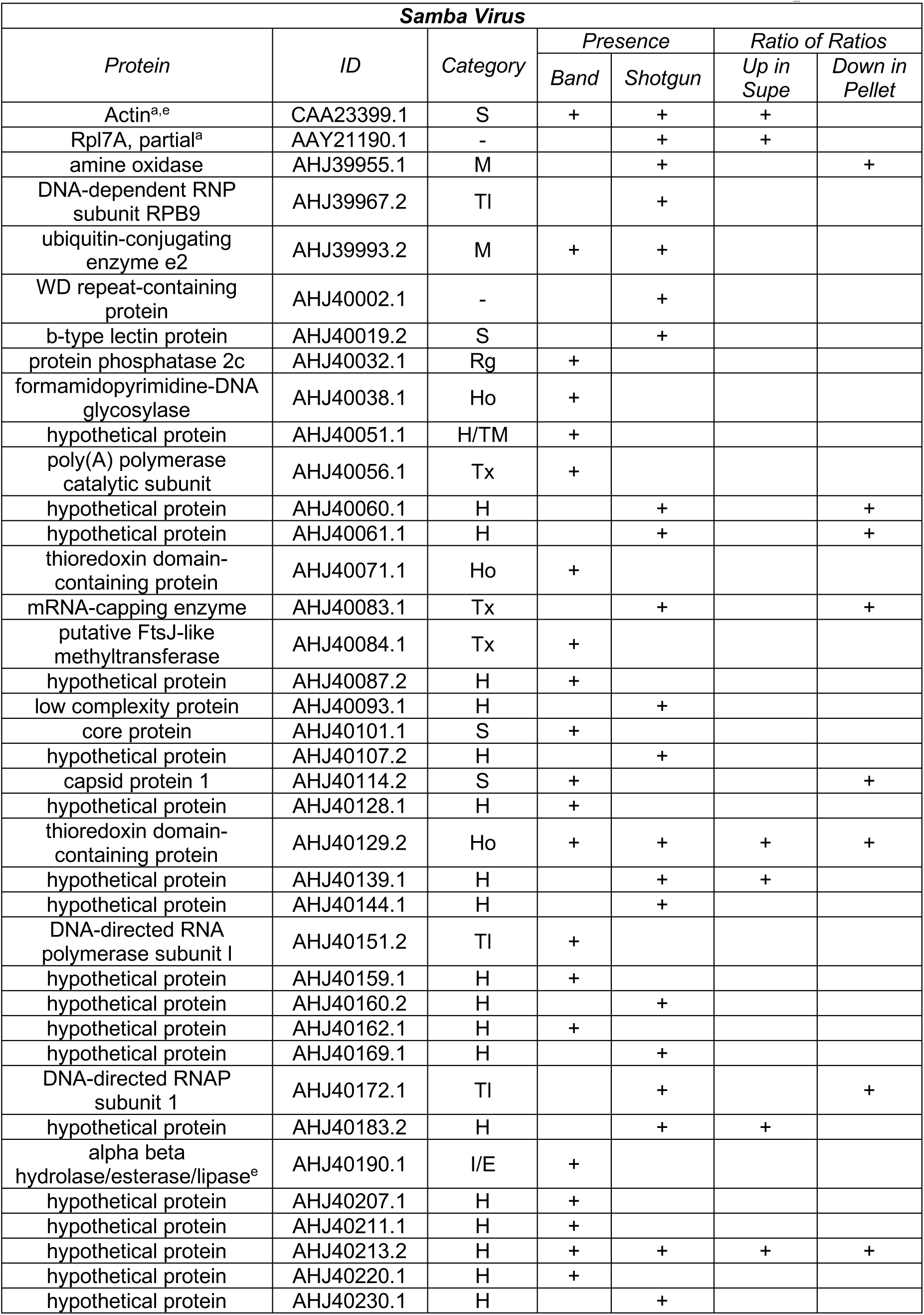
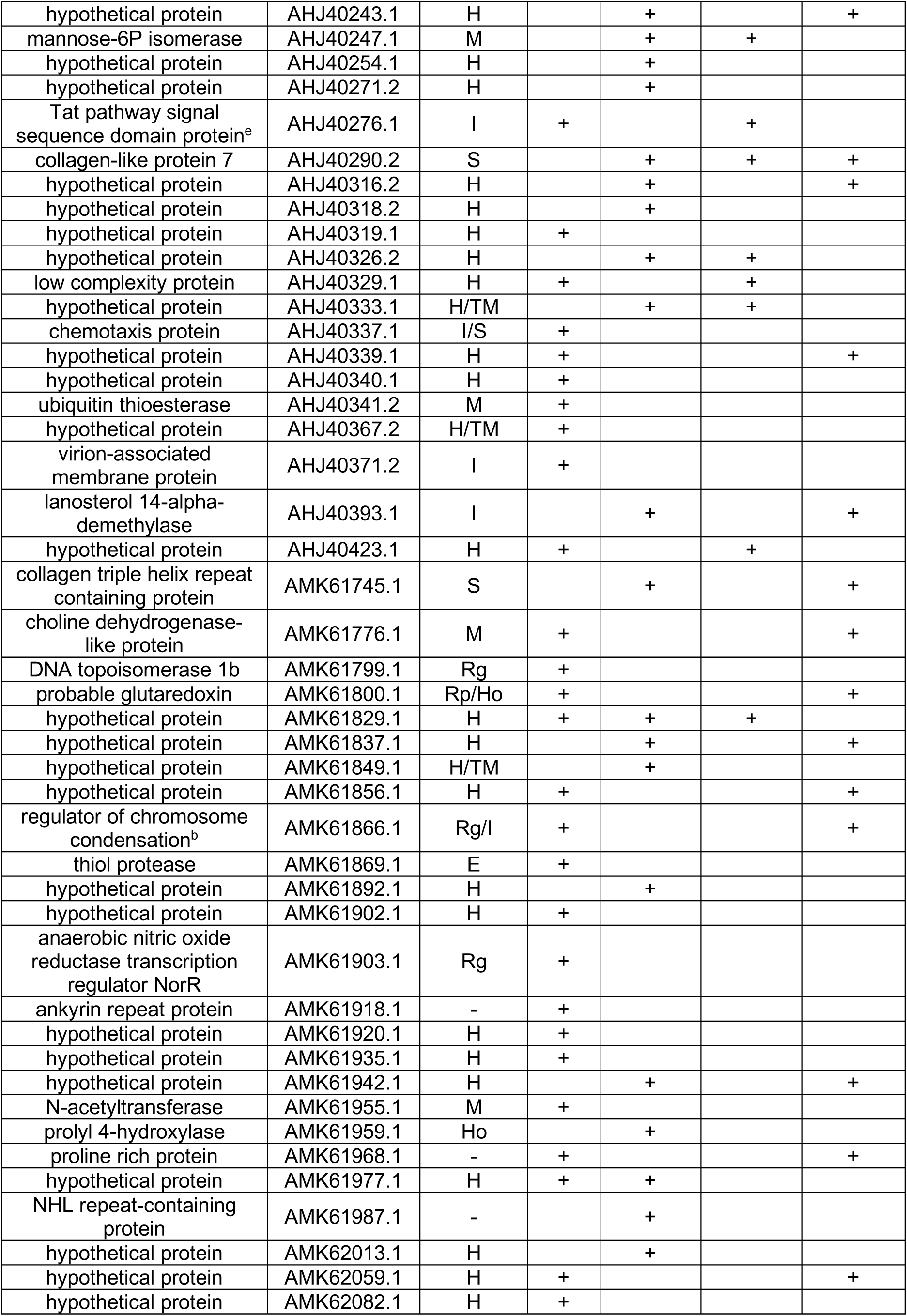
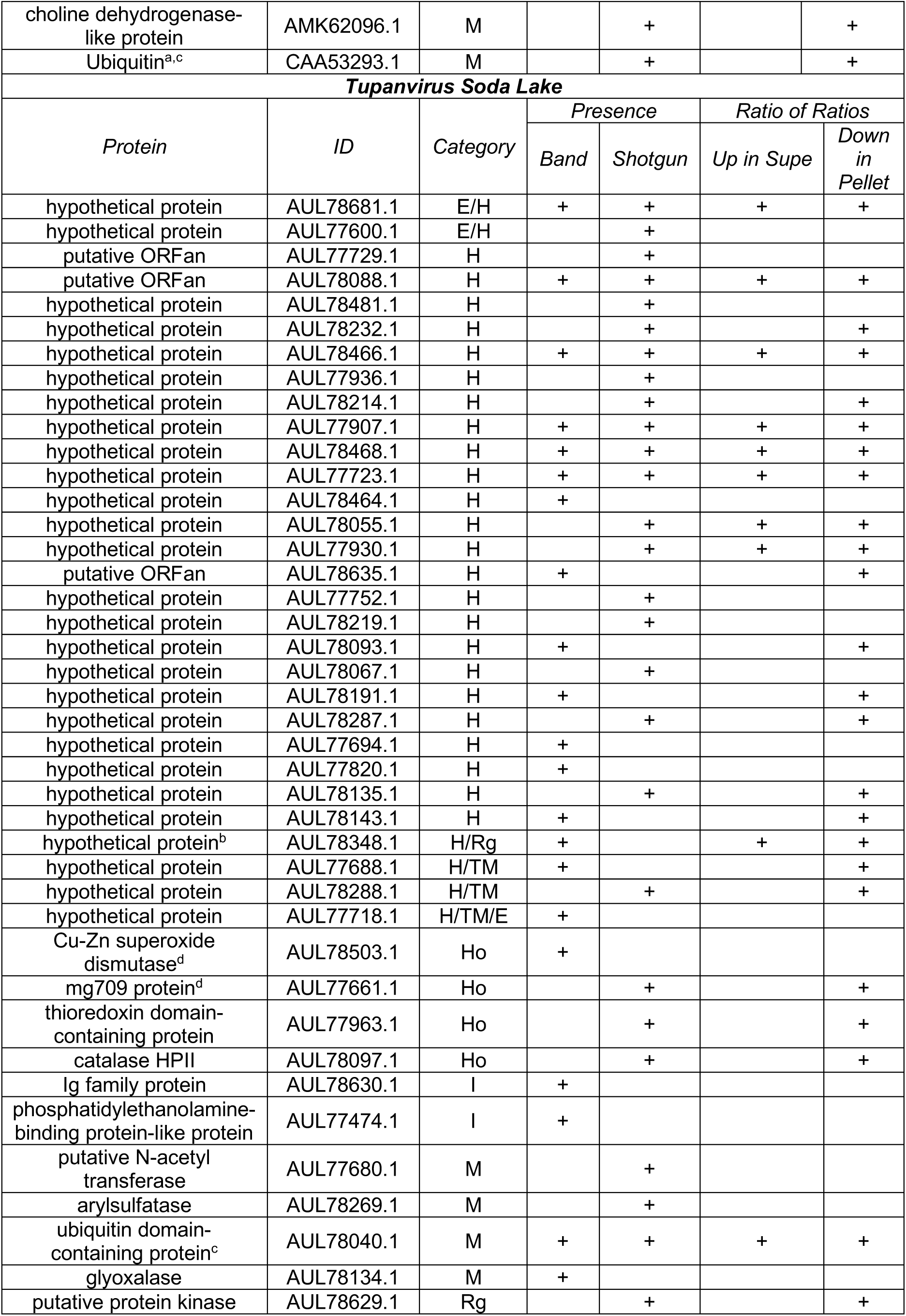
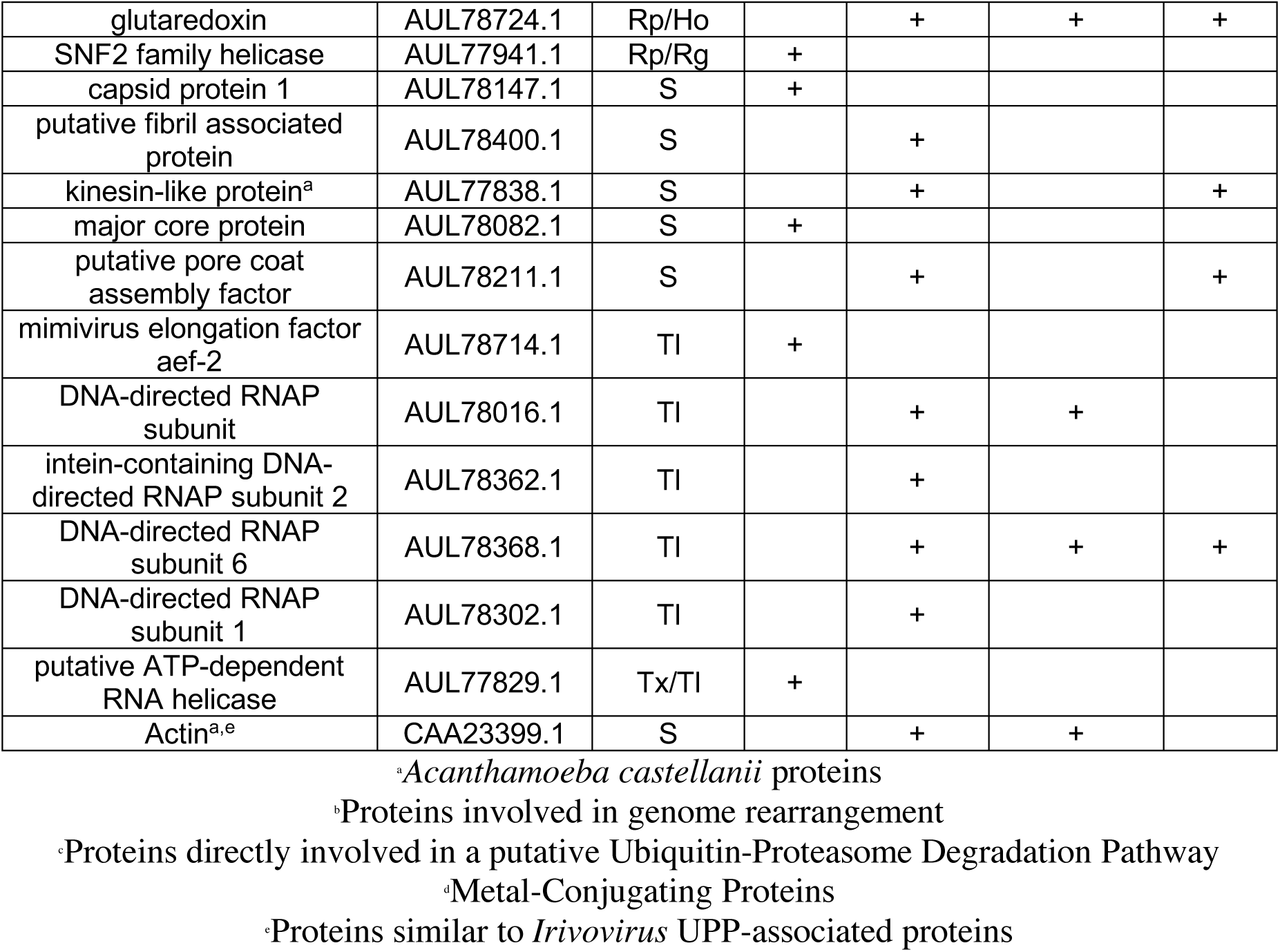
Identification of Proteins Released from SMBV and TV Capsids.

Additionally, we have identified proteins that are released during the initial stages of infection in two GV, SMBV and TV. Over half of the proteins released by these viruses are annotated as hypothetical, low complexity, or as an ORFan. We were able to provide functional predictions for some of these proteins through homology. Even so, an exact functional determination of these proteins remains elusive. The release of these proteins at the initiation of stargate opening suggests that these proteins play an important role in the early stages of GV infection (phagosome survival, genome translocation, early transcription, host defense suppression, etc.). The exact functions of these proteins, as well as how their interactions mediate and orchestrate GV infection, are prime candidates for future study. The importance of these potential future studies is enhanced by the fact that many GV appear to share similar strategies for genome release. All four of the GV tested in this study responded to the treatment conditions, suggesting that these GV utilize similar molecular forces during genome release, and likely similar proteins to counteract these forces.

## Abbreviations

SMBV: Samba virus
TV: Tupanvirus soda lake
GV: Giant virus
POP: Percentage of open particles
UPP: Ubiquitin-proteasome degradation pathway

## Acknowledgements

The authors would like to thank the MSU Proteomics and Mass Spec. Facilities, especially Drs. D. Jones, C. Wilkerson, and D. Whiten, for their assistance with MS experiments. Additionally, we thank Drs. W. Jiang, T. Klose, and V. Bowman at Purdue University’s Midwest Cryo-EM Consortium (NIH Consortium #U24GM116789-03). We would also like to thank C. Flegler at the MSU Center for Advanced Microscopy for her expertise with SEM experiments. The MSU High Performance Computation Cluster (HPCC) provided computational tools and support for cryo-EM image motion correction. Dr. K. Padmanabhan and Dr. M. Feig provided additional assistance with computational resources and expert consultation in protein homology and functional predictions. Funding for this project was provided by the AAAS Marion Mason Milligan award for Women in the Chemical Sciences (KNP), the JK Billman, Jr., MD Endowed Research Professorhip (KNP), Funding was provided by Fundo de Amparo à Pesquisa do Estado do Rio de Janeiro e Conselho Nacional de Pesquisa (JRC), and the Burroughs Wellcome Fund (KNP). JRS has been supported by the Jack Throck Waston Fellowship and the August and Ernest Frey Research Fellowship from MSU and NIH R01 GM110185 (KNP). JSA has been supported by CNPq, CAPES, MS, and FAPEMIG. Nividia provided GPU support for cryo-EM and cryo-ET image processing.

## Author Contributions

JRS: Designed and carried out experiments, wrote the manuscript, data analysis

JRC: Viral sample preparation and manuscript review, data analysis

JSA: manuscript review

KNP: Designed and carried out experiments, manuscript review, data analysis

## STAR Methods

### Contact for Reagent and Resource Sharing

Further information and requests for resources and reagents should be directed to and will be fulfilled by the Lead Contact, Kristin Parent.

### Experimental Model and Subject Details

#### Acanthamoeba castellanii

*Acanthamoeba castellanii* cells were purchased from ATCC (ATCC 30010). *Acanthamoeba castellanii* (ATCC 30010) was cultivated in 712 PYG media w/Additives (ATCC recipe) at pH 6.5 in the presence of gentamicin (15 μg/mL) and penicillin/streptomycin (100 U/mL) at 28 °C to reach a 90% confluence.

#### Giant Viruses

Tupanvirus soda lake (TV), Antarctica virus, and Samba virus (SMBV) were isolated previously (Abrahao et al., 2018; Andrade et al., 2018; Campos et al., 2014). M4 virus was kindly provided by Dr. Bernard La Scola and Dr. Thomas Klose (Boyer et al., 2011). *Acanthamoeba castellanii* (ATCC 30010) was cultivated in 712 PYG media w/ Additives (ATCC recipe) at pH 6.5 in the presence of gentamicin (15 μg/mL) and penicillin/streptomycin (100 U/mL) at 28 °C to reach a 90% confluence. SMBV or TV virions were diluted in phosphate buffered saline (PBS) and added to the cells to a multiplicity of infection of 5 (TV) or 10 (SMBV). An initial incubation was carried out for one hour at room temperature. After the initial incubation, additional PYG media was added to the cells and the flasks were incubated at 28 °C for 48 hours. After 48 hours, more of the free amoebal cells had been lysed. Suspensions containing cell debris and cell particles were centrifuged at 900 x g to pellet residual cells. The resulting supernatant was filtered using a 2 μm filter and was immediately applied to a 22% sucrose cushion (w/w) at 15,000 x g for 30 min. Viral pellets were resuspended in PBS and stored at -80 °C. Viruses were tittered using the Reed-Muench protocol (Ramakrishnan, 2016). On average, virus isolation yielded 10^10^ TCID_50_ /mL (TCID = tissue culture infective dose).

### Method Details

#### Treatment of SMBV Particles and Image Analysis

##### Determining the Percentage of Open SMBV Particles

For all treatments, the percentage of open SMBV particles (POP) was determined via single particle cryo-electron microscopy. These percentages were compared to the native (untreated) level of spontaneous SMBV particle opening, determined previously to be ∼5% (Schrad et al., 2017).

##### Conditions That Did Not Increase POP

SMBV particles were treated with various conditions that have been shown to disrupt/destroy other viruses. These conditions include urea, guanidinium hydrochloride, DMSO, Triton X-100, chloroform, DNase I, and an enzyme cocktail (lysozyme, bromelain, proteinase K) that was previously shown to remove APMV fibers (Kuznetsov et al., 2010). Treatments were applied for 1-2 hours prior to POP determination via cryo-EM. Concentrations for the various conditions, as well as the resultant POP values, can be found in Table S1.

##### pH Titration of SMBV Particles

25-50 μL of SMBV particles were added to Millipore VSWP Membrane Filter dialysis discs (0.025 μm cutoff) which were then floated onto ∼25 mL of 20 mM sodium phosphate buffer, adjusted to the desired pH. The samples were allowed to equilibrate for 1.5-2 hours. For conditions where low pH would interfere with additional treatment (*e.g.* pH 2 + DNase I or pH 2 samples submitted for mass spectrometry) the particles were dialyzed for an additional 1.5-2 hours against pH 7.0 buffer to restore neutral pH.

##### High Temperature Incubation

GV particles were incubated in a BioRad T100 thermal cycler at 80, 89, and 100 °C for 1 hour. SMBV particles remained intact following 1 hours at 100 °C, so additional incubations at 100 °C were performed at 2, 3, or 6 hours. As a control, SMBV particles were also incubated for 1 hour at room temperature (25 °C).

##### Combining High Temperature and Low pH

To determine the effect of combining low pH and high temperature, GV particles were sequentially treated with pH 2 and 100 °C. First, SMBV particles were dialyzed against 20 mM sodium phosphate buffer, adjusted to pH 2, for 2 hours. Following dialysis, SMBV particles were incubated at 100 °C for 3 hours.

#### Cryo-Electron Microscopy (Cryo-EM) and Cryo-Electron Tomography (Cryo-ET)

##### Sample Preparation

Samples for cryo-EM and cryo-ET were prepared as described previously (Schrad et al., 2017). Briefly, small (3-5 μL) aliquots of virus particles were applied to R2/2 (cryo-EM) or R 3.5/1 (cryo-ET) Quantifoil grids (Electron Microscopy Solutions) that had been plasma cleaned for 20 seconds in a Fischione model 1020 plasma cleaner. Prior to virus addition, 5-10 μL of 10 nm nanogold fiducial markers were applied to the R3.5/1 grids and were air dried to provide markers for fiducial alignment of the tilt series. The samples were plunge frozen in liquid ethane using a manual plunge-freezing device (Michigan State University Physics Machine Shop). Frozen-hydrated samples were stored, transferred, and imaged under liquid nitrogen temperatures.

##### Single Particle Cryo-Electron Microscopy

Single particle cryo-EM experiments were performed at Michigan State University. Virus particles were imaged in a JEOL 2200-FS TEM operating at 200 keV, using low dose conditions controlled by SerialEM (version 3.5.0-beta, (Mastronarde, 2005)) with the use of an in-column Omega Energy Filter operating at a slit width of 35 eV. Micrographs were recorded at 25 frames per second using a Direct Electron DE-20 direct detector, cooled to -38 °C. Motion correction was performed using the Direct Electron software package (Direct Electron, LLC). Micrographs were collected between 8,000 and 10,000 X nominal magnification (6.87 and 5.30 Å/pixel, respectively). The objective lens defocus settings ranged from 10 to 15 μm underfocus. Micrographs were collected for 5 seconds, resulting in a total dose of ∼35 e-/Å^2^. For bubblegram imaging, the SMBV particles were imaged for an additional four exposures, resulting in a total does of ∼140 e-/Å^2^

##### Cryo-Electron Tomography

Cryo-ET tilt series were collected using a Titan Krios TEM operating at 300 keV with a post-column GIF (20 eV slit width) under low dose conditions controlled by SerialEM or Leginon at Purdue University. Images were collected using a Gatan K2 direct electron detector operating at 100 milliseconds/frame. Images were collected in super resolution mode between 33,000 and 53,000 X nominal magnification (2.12 - 1.33 Å/pixel). Tilt series were carried out between +/-50 ° with bidirectional image collection every 2 °. Images were collected for 5 seconds, resulting in ∼2.5 electrons/Å^2^ per tilt image (∼125 electrons/Å^2^, total exposure dose).

Individual micrographs were corrected for particle motion and binned by a factor of two using MotionCor2 v1.2.0 (Zheng et al., 2017) and the corrected images were stitched back into a tilt series using the newstack functionality in IMOD (Mastronarde and Held, 2017). Tilt series alignment, using fiducial markers, and tomogram generation was carried out using IMOD v4.7.5. Final tomogram volumes were generated using ten iterations of the SIRT reconstruction method (Mastronarde, 1997) then filtered using the smooth (3×3 kernel) and median (size 3) options in IMOD. Select tomograms were annotated using Amira v2019.2 (ThermoFisher Scientific).

#### Scanning Electron Microscopy

##### SEM Preparation and Imaging

GV particles were imaged using a JEOL JSM-7500F scanning electron microscope. Prior to imaging, virus particles were desiccated using an EM CPD300 critical point dryer, fixed with glutaraldehyde onto poly-L-Lysine treated SEM slides, and sputter coated with a ∼2.7nm layer of iridium using a Q150T Turbo Pumped Coater. Particles were imaged between 8,500 X and 85,000 X nominal magnification.

#### Differential Mass Spectrometry

##### Sample Preparation

SMBV and TV particles were dialyzed against 20 mM sodium phosphate buffer, adjusted to pH 2, for 2 hours, as described above. An aliquot of each virus was left undialyzed as a control (Material Applied, MA). Following dialysis, proteins that had been released from the viral particles were separated from the virions via centrifugation in a microcentrifuge at 8,000 x g for 15 minutes. Visible viral pellets were resuspended in the same volume as the supernatant using 20 mM sodium phosphate buffer, adjusted to pH 7.0. Two technical replicates were created for each sample. An aliquot of each sample was used for SDS-PAGE.

Each sample was TCA precipitated and submitted for LC/MS/MS analysis to the MSU Proteomics Core. Prior to submission, samples were run on a 15% polyacrylamide gel at a voltage of 200 V for 45 minutes. TV and SMBV gel bands visible by Coomassie blue stain were excised and submitted for MS analysis as well.

##### Proteolytic Digestion

TCA precipitated pellets were re-suspended in 270uL of 100mM ammonium bicarbonate supplemented with 10% trifluoroethanol. Samples were reduced and alkylated by adding TCEP and Iodoacetamide at 10mM and 40mM, respectively and incubating for 5min at 45C with shaking at 1400 rpm in an Eppendorf ThermoMixer. Trypsin, in 100mM ammonium bicarbonate, was added at a 1:100 ratio (wt/wt) and the mixture was incubated at 37C overnight. Final volume of each digest was ∼300uL. After digestion, the samples were acidified to 2% TFA and subjected to C18 solid phase clean up using StageTips(Rappsilber et al., 2007) to remove salts.

##### LC/MS/MS and Data Analysis

An injection of 5uL was automatically made using a Thermo EASYnLC 1200 onto a Thermo Acclaim PepMap RSLC 0.075mm x 20mm C18 trapping column and washed for ∼5min with buffer A. Bound peptides were then eluted over 95min with a gradient of 8%B to 42%B in 84min, ramping to 100%B at 85min and held at 100%B for the duration of the run (Buffer A = 99.9% Water/0.1% Formic Acid, Buffer B = 80% Acetonitrile/0.1% Formic Acid/19.9% Water) at a constant flow rate of 300nl/min. Column temperature was maintained at a constant temperature of 50 °C using and integrated column oven (PRSO-V2, Sonation GmbH, Biberach, Germany). Eluted peptides were sprayed into a ThermoScientific Q-Exactive HF-X mass spectrometer using a FlexSpray spray ion source. Survey scans were taken in the Orbi trap (60,000 resolution, determined at m/z 200) and the top ten ions in each survey scan are then subjected to automatic higher energy collision induced dissociation (HCD) with fragment spectra acquired at 7,500 resolution. The resulting MS/MS spectra are converted to peak lists using MaxQuant v1.6.0.1(Cox and Mann, 2008) and searched using the Andromeda (Cox et al., 2011) algorithm against a protein database containing sequences from SMBV or TV and *Acanthamoeba castellanii* (each downloaded from NCBI, www.ncbi.nlm.nih.gov). Common laboratory contaminants were included in the Andromeda search. Protein and peptide FDR for all searches were set to 1%.

##### Mass Spectrometry Data Synthesis

The percentage of the total LFQ signal each protein was responsible for in each sample was calculated by dividing the individual protein LFQ signal by the total LFQ signal for the sample, excluding contaminates. Proteins that are released from the viral particles are expected to make up a higher percentage of the supernatant sample than the whole virion (MA), so the ratios of these two percentages were calculated (Table 2, Table S3). Proteins with a supernatant:MA ratio > 1 were selected for further analysis.

##### Classification/Functional Annotation of Proteins Identified via MS

TV and SMBV proteins released at low pH were classified via their predicted functions and domains. Primary functional annotation had been carried out previously for both TV (Abrahao et al., 2018) and SMBV (Campos et al., 2014). Additional functional prediction, as well as homology prediction between the two viruses, was carried out through the use of the NCBI BLAST database (NCBI) as well as the HHBLITS server (Remmert et al., 2011) and the InterPro database (Mitchell et al., 2019). Domain prediction was carried out by searching the InterPro database and utilizing the PSIPRED server (Buchan and Jones, 2019) with the DISOPRED3 (Jones and Cozzetto, 2015) functionality activated.

### Quantification and Statistical Analysis

#### Mass Spectrometry Analysis

LFQ intensities for both SMBV and TV spectra were detected in triplicate. For each virus, the initial run did not produce high quality data so these intensities were disregarded. LFQ intensities from the remaining two runs were averaged together to produce the reported intensity (Table S3).

### Data and Software Availability

Three-dimensional tomograms have been deposited to the Electron Microscopy Database (EMDB) under the ID codes EMD-20747 (pH 2, Movie S3), EMD-20748 (100 °C, Movie S4), EMD-20746 (pH 2 + 100 °C, Movie S6), and EMD-20745 (pH 2 + 100 °C, Movie S8).

## Supplemental File Legends

Supplemental Movie 1: **Untreated SMBV Bubblegram Imaging. Related to Figure 1.** Bubblegram image series of a native SMBV particle demonstrating the buildup of radiation damage over time. A clear star-shaped radiation damage pattern is observed around the 11:00 position on the particle. Each frame represents a two second exposure (14 e-/Å^2^). Total exposure time = 24 seconds (∼140 e-/Å^2^).

Supplemental Movie 2: **Untreated SMBV Tomogram. Related to Figure 2.** Slice-by-slice view of a tomogram of a native SMBV particle. Depicted in Figure 2B-C within the main body of the manuscript.

Supplemental Movie 3: **Low pH Treated SMBV Tomogram. Related to Figure 2.** Slice-by-slice view of a tomogram of a pH 2-treated SMBV particle. Note the opening in the stargate vertex as well as the sac exiting the capsid. Depicted in Figure 2F-G.

Supplemental Movie 4: **Tomogram of SMBV Incubated at High Temperature. Related to Figure 2.** Slice-by-slice view of a tomogram from an SMBV particle incubated at 100 °C for 6 hours. Note the fully open stargate vertex, the exodus of the nucleocapsid, and the apparent tethers between the capsid and the nucleocapsid. Depicted in Figure 2J-K.

Supplemental Movie 5: **Tilt Series of High Temperature Incubated SMBV. Related to Figure 2.** Tilt series of an SMBV particle incubated at 100 °C. Tilts were acquired every 2 degrees ranging from +/-## degrees. Depicted in Figure 2J-K.

Supplemental Movie 6: **Low pH and High Temperature Treated SMBV Tomogram. Related to Figure 2.** Slice-by-slice view of a tomogram of an SMBV particle treated with both low pH and high temperature. Tomogram segmentation was carried out using Amira v2019.2. Colors represent the following: Red-Outer Capsid Layer, Orange-Inner Capsid Layer, Blue-Starfish Seal Complex, and Yellow-Lipid. Note the flexibility of the innermost capsid layer and the residual density within the capsid interior. Depicted in Figure 2N-O.

Supplemental Movie 7: **Low pH and High Temperature Treated SMBV Tilt Series. Related to Figure 2.** Tilt series of an SMBV particle treated with both pH 2 and 100 °C. Tilts were acquired every 2° ranging from +/-50°degrees. Depicted in Figure 2N-O.

Supplemental Movie 8: **Low pH and High Temperature Treated SMBV Tomogram. Related to Figure 2.** Slice-by-slice view of a tomogram of five SMBV particles treated with both low pH and high temperature. These particles all have open stargate vertices, and one is oriented in a top-down view, providing additional structural information about the SMBV particle.

Supplemental Movie 9: **Low pH and High Temperature Treated SMBV Tilt Series. Related to Figure 2.** Tilt series of an SMBV particle treated with both pH 2 and 100 °C. Tilts were acquired every 2° ranging from +/-50°degrees. Five distinct SMBV particles are visible within this tilt series.

Supplemental Table 1: **Conditions That SMBV Particles Resist. Related to Figure 1.** Treatment conditions that did not produce a marked increase in the percentage of open SMBV particles.

Supplemental Table 2: **SMBV and TV Released Hypothetical Proteins With Predicted Functionalities. Related to Figure 5.** Functional predictions of SMBV and TV hypothetical proteins from BLAST, HHBLITS, PSIPRED, or InterPro Analysis.

Supplemental Table 3: SMBV and TV Proteins With LFQ Percentages and Comparison Between Supernatant and Pellet Levels. Related to Figure 5. Table of the proteins identified through the shotgun mass spectrometry experiments for SMBV and TV. Percentages for the Material Applied, Pellet, and Supernatant samples represent the percentage of the overall signal that each protein accounted for in the LFQ intensities. Supernatant/MA and Pellet/MA values represent the relative contribution of each protein to the given sample’s spectral intensity as compared to the untreated particles (MA).

Supplemental Figure 1: **Percentage of Fiberless SMBV Particles at Varying Temperatures. Related to Figure 1.** Histogram of the percentage of fiberless (open or unopen) SMBV particles at various temperatures and incubation times.

Supplemental Figure 2: **Sample Preparation for SDS-PAGE and LC/MS/MS Experiments. Related to Figures 4 & 5.** A cartoon of the workflow schematic used to prepare samples for both the SDS-PAGE and LC/MS/MS experiments.

