## Supplemental Table 1 for "Boiling Acid Mimics Intracellular Giant Virus Genome Release"

Table S1

| **Condition** | **Concentration(s)** | **% Open SMBV** |
| --- | --- | --- |
| Urea | 9M | 2.94 |
| Guanidinium Hydrochloride | 3M, 6M | 2.90 |
| DMSO | 1% (v/v) | 2.00 |
| Triton X-100 | 1% (v/v) | 0.00 |
| Bromelain, Proteinase K, Lysozyme | 14, 1, 10 mg/mL | 0.00 |
| Chloroform | 20% (v/v) | 4.17 |
| DNase I | 2 mg/mL | 2.33 |
