## Supplemental Table 2 for "Boiling Acid Mimics Intracellular Giant Virus Genome Release"

| ***Virus*** | ***Accession Number*** | **Functional Prediction** | **Functional Prediction** |
| --- | --- | --- | --- |
| *SMBV* | *AHJ4005.1.1* | TM Helix |  |
| *SMBV* | *AHJ40159.1* | TM Helix |  |
| *SMBV* | *AHJ40213.2* | Methyl-Accepting Chemotaxis Protein |  |
| *SMBV* | *AHJ40326.1* | Alpha L Rhamnidose Domain |  |
| *SMBV* | *AHJ40333.1* | Beta galactosidase Domain | SpoIID/LytB Domain |
| *SMBV* | *AHJ40333.1* | TM Helix |  |
| *SMBV* | *AHJ40367.2* | TM Helix | Apple Domain (Proteolysis) |
| *SMBV* | *AHJ40423.1* | Crp/Fnr Family Transcription Regulator |  |
| *SMBV* | *AMK61829.1* | Coiled-Coil Domain-Containing Protein 180-like Isoform |  |
| *SMBV* | *AMK61849.1* | TM Helix |  |
| *SMBV* | *AMK61920.1* | Ankyrin Repeat Protein |  |
| *SMBV* | *AMK61942.1* | Alpha/Beta Hydrolase | Zn-Finger Protein |
| *SMBV* | *AMK61942.1* | TM Helix |  |
| *SMBV* | *AMK62013.1* | Coiled-Coil and C2 Domain-Containing Protein 1-like Isoform |  |
| *SMBV* | *AMK62059.1* | LamG Superfamily (Incomplete Domain) |  |
| *TV* | *AUL77600.1* | Alpha/Beta Hydrolase |  |
| *TV* | *AUL77688.1* | TM Helix |  |
| *TV* | *AUL77718.1* | Apple Domain (Proteolysis) | TM Helix |
| *TV* | *AUL77723.1* | Membrane Helix |  |
| *TV* | *AUL77930.1* | Sugar O-Acetyltransferase | Outer Dense Fiber Protein 3-B-like Protein |
| *TV* | *AUL78055.1* | ATPase | Ion Channel |
| *TV* | *AUL78135.1* | (Ribo)Nuclease Hydrolase | Inosine-uridine Nucleoside Hydrolase |
| *TV* | *AUL78143.1* | Nuclear Transport Family 2 Protein | Sgc/EcaC Family Oxidoreductase |
| *TV* | *AUL78288.1* | TM Helix |  |
| *TV* | *AUL78348.1* | MC1 Domain (DNA Protection) | Ubiquitin-Conjugating Enzyme |
| *TV* | *AUL78681.1* | Apple Domain (Proteolysis) |  |
