## Supplemental Table 3 for "Boiling Acid Mimics Intracellular Giant Virus Genome Release"

| ***Samba virus*** | | | | | | | | | | | | | | | | | |
| --- | --- | --- | --- | --- | --- | --- | --- | --- | --- | --- | --- | --- | --- | --- | --- | --- | --- |
| *Protein* | *ID* | *Pep* | *Material Applied* | | | *Pellet* | | | *Supernatant* | | | *Supernatant/MA* | | | *Pellet/MA* | | |
|  |  |  | *2* | *3* | *Avg.* | *2* | *3* | *Avg.* | *2* | *3* | *Avg.* | *2* | *3* | *Avg.* | *2* | *3* | *Avg.* |
|  |  |  | *%* | *%* | *%* | *%* | *%* | *%* | *%* | *%* | *%* |  |  |  |  |  |  |
| actin | CAA23399.1 | 11 | 0.26 | 0.28 | 0.27 | 1.02 | 0.26 | 0.64 | 1.69 | 0.91 | 1.30 | 6.48 | 3.26 | 4.87 | 3.90 | 0.94 | 4.84 |
| Rpl7A, partial | AAY21190.1 | 2 | 0.03 | 0.05 | 0.04 | 0.05 | 0.02 | 0.04 | 0.12 | 0.25 | 0.18 | 3.68 | 5.46 | 4.57 | 1.52 | 0.48 | 1.99 |
| mannose-6P isomerase | AHJ40247.1 | 10 | 0.08 | 0.01 | 0.05 | 0.33 | 0.13 | 0.23 | 0.03 | 0.08 | 0.06 | 0.41 | 7.89 | 4.15 | 3.99 | 12.92 | 16.91 |
| Tat pathway signal sequence domain protein | AHJ40276.1 | 9 | 0.09 | 0.06 | 0.07 | 0.20 | 0.37 | 0.28 | 0.20 | 0.27 | 0.23 | 2.13 | 4.68 | 3.41 | 2.15 | 6.36 | 8.51 |
| hypothetical protein | AMK62013.1 | 8 | 0.13 | 0.10 | 0.12 | 0.00 | 0.23 | 0.11 | 0.00 | 0.58 | 0.29 | 0.00 | 5.90 | 2.95 | 0.00 | 2.33 | 2.33 |
| collagen-like protein 7 | AHJ40290.2 | 6 | 0.20 | 0.10 | 0.15 | 0.00 | 0.04 | 0.02 | 0.15 | 0.31 | 0.23 | 0.75 | 3.21 | 1.98 | 0.00 | 0.38 | 0.38 |
| hypothetical protein | AMK61829.1 | 25 | 1.73 | 0.33 | 1.03 | 0.74 | 1.16 | 0.95 | 0.45 | 1.08 | 0.76 | 0.26 | 3.28 | 1.77 | 0.43 | 3.55 | 3.97 |
| hypothetical protein | AHJ40139.1 | 25 | 0.09 | 0.03 | 0.06 | 0.08 | 0.22 | 0.15 | 0.00 | 0.11 | 0.06 | 0.00 | 3.21 | 1.60 | 0.98 | 6.38 | 7.36 |
| hypothetical protein | AHJ40423.1 | 5 | 0.09 | 0.10 | 0.09 | 0.09 | 0.12 | 0.11 | 0.11 | 0.17 | 0.14 | 1.23 | 1.66 | 1.45 | 1.07 | 1.18 | 2.25 |
| hypothetical protein | AHJ40333.1 | 7 | 0.84 | 0.78 | 0.81 | 0.60 | 2.28 | 1.44 | 0.54 | 1.54 | 1.04 | 0.64 | 1.98 | 1.31 | 0.72 | 2.94 | 3.66 |
| hypothetical protein | AHJ40183.2 | 11 | 0.03 | 0.00 | 0.01 | 0.00 | 0.02 | 0.01 | 0.00 | 0.01 | 0.01 | 0.00 | 2.33 | 1.16 | 0.00 | 4.49 | 4.49 |
| hypothetical protein | AHJ40326.2 | 4 | 0.26 | 0.24 | 0.25 | 0.52 | 0.45 | 0.48 | 0.32 | 0.26 | 0.29 | 1.21 | 1.07 | 1.14 | 1.97 | 1.87 | 3.83 |
| thioredoxin domain-containing protein | AHJ40129.2 | 10 | 0.68 | 0.66 | 0.67 | 0.27 | 0.58 | 0.42 | 0.00 | 1.35 | 0.67 | 0.00 | 2.02 | 1.01 | 0.40 | 0.87 | 1.26 |
| low complexity protein | AHJ40329.1 | 22 | 16.10 | 18.2 | 17.17 | 56.21 | 28.85 | 42.53 | 19.39 | 14.38 | 16.89 | 1.20 | 0.79 | 1.00 | 3.49 | 1.58 | 5.07 |
| Ubiquitin-60S ribosomal protein L40 | CAA53293.1 | 6 | 0.21 | 0.27 | 0.24 | 0.10 | 0.10 | 0.10 | 0.20 | 0.26 | 0.23 | 0.93 | 0.97 | 0.95 | 0.45 | 0.36 | 0.82 |
| hypothetical protein | AHJ40230.1 | 5 | 0.26 | 0.37 | 0.32 | 0.55 | 0.54 | 0.55 | 0.30 | 0.25 | 0.28 | 1.16 | 0.68 | 0.92 | 2.08 | 1.48 | 3.56 |
| probable glutaredoxin | AMK61800.1 | 5 | 0.04 | 0.05 | 0.05 | 0.00 | 0.04 | 0.02 | 0.00 | 0.10 | 0.05 | 0.00 | 1.77 | 0.89 | 0.00 | 0.72 | 0.72 |
| ubiquitin-conjugating enzyme e2 | AHJ39993.2 | 6 | 1.03 | 1.10 | 1.07 | 1.32 | 1.03 | 1.17 | 0.79 | 0.87 | 0.83 | 0.76 | 0.80 | 0.78 | 1.27 | 0.94 | 2.21 |
| proline rich protein | AMK61968.1 | 13 | 0.86 | 0.93 | 0.89 | 0.29 | 0.97 | 0.63 | 0.35 | 1.03 | 0.69 | 0.41 | 1.11 | 0.76 | 0.34 | 1.04 | 1.38 |
| hypothetical protein | AHJ40087.2 | 13 | 0.69 | 0.26 | 0.47 | 0.54 | 1.43 | 0.99 | 0.00 | 0.37 | 0.18 | 0.00 | 1.44 | 0.72 | 0.78 | 5.61 | 6.39 |
| lanosterol 14-alpha-demethylase | AHJ40393.1 | 3 | 0.07 | 0.43 | 0.25 | 0.09 | 0.27 | 0.18 | 0.08 | 0.09 | 0.09 | 1.11 | 0.22 | 0.66 | 1.21 | 0.62 | 1.83 |
| b-type lectin protein | AHJ40019.2 | 4 | 0.01 | 0.01 | 0.01 | 0.02 | 0.08 | 0.05 | 0.00 | 0.02 | 0.01 | 0.00 | 1.27 | 0.63 | 1.21 | 6.29 | 7.51 |
| thioredoxin domain-containing protein | AHJ40071.1 | 27 | 1.72 | 0.88 | 1.30 | 0.91 | 1.44 | 1.18 | 0.25 | 0.91 | 0.58 | 0.15 | 1.03 | 0.59 | 0.53 | 1.64 | 2.17 |
| hypothetical protein | AHJ40169.1 | 3 | 0.92 | 0.74 | 0.83 | 0.91 | 1.37 | 1.14 | 0.36 | 0.55 | 0.46 | 0.39 | 0.75 | 0.57 | 0.99 | 1.85 | 2.84 |
| kinesin-like protein | AHJ40024.1 | 44 | 0.10 | 0.03 | 0.06 | 0.00 | 0.31 | 0.15 | 0.00 | 0.03 | 0.02 | 0.00 | 1.12 | 0.56 | 0.00 | 11.15 | 11.15 |
| low complexity protein | AHJ40093.1 | 13 | 1.52 | 0.96 | 1.24 | 1.31 | 1.42 | 1.36 | 0.33 | 0.85 | 0.59 | 0.22 | 0.89 | 0.55 | 0.87 | 1.47 | 2.34 |
| hypothetical protein | AHJ40162.1 | 11 | 1.42 | 1.69 | 1.55 | 0.85 | 0.85 | 0.85 | 0.88 | 0.82 | 0.85 | 0.62 | 0.49 | 0.55 | 0.60 | 0.50 | 1.10 |
| hypothetical protein | AHJ40160.2 | 11 | 0.39 | 0.25 | 0.32 | 0.37 | 0.47 | 0.42 | 0.00 | 0.26 | 0.13 | 0.00 | 1.03 | 0.52 | 0.96 | 1.87 | 2.82 |
| hypothetical protein | AHJ40213.2 | 24 | 1.00 | 1.05 | 1.02 | 0.70 | 0.86 | 0.78 | 0.45 | 0.59 | 0.52 | 0.45 | 0.57 | 0.51 | 0.70 | 0.83 | 1.52 |
| anaerobic nitric oxide reductase transcription regulator NorR | AMK61903.1 | 7 | 0.29 | 0.21 | 0.25 | 0.19 | 0.42 | 0.30 | 0.05 | 0.14 | 0.09 | 0.17 | 0.66 | 0.41 | 0.67 | 1.97 | 2.64 |
| core protein | AHJ40101.1 | 44 | 1.19 | 0.50 | 0.85 | 0.45 | 1.43 | 0.94 | 0.21 | 0.32 | 0.27 | 0.18 | 0.64 | 0.41 | 0.38 | 2.87 | 3.25 |
| ubiquitin thioesterase | AHJ40341.2 | 10 | 0.53 | 0.49 | 0.51 | 0.32 | 0.28 | 0.30 | 0.17 | 0.21 | 0.19 | 0.32 | 0.42 | 0.37 | 0.60 | 0.57 | 1.17 |
| hypothetical protein | AMK61920.1 | 30 | 3.61 | 2.70 | 3.15 | 1.10 | 2.08 | 1.59 | 0.33 | 1.69 | 1.01 | 0.09 | 0.63 | 0.36 | 0.30 | 0.77 | 1.07 |
| hypothetical protein | AHJ40271.2 | 8 | 0.31 | 0.41 | 0.36 | 0.31 | 1.37 | 0.84 | 0.00 | 0.28 | 0.14 | 0.00 | 0.69 | 0.34 | 1.02 | 3.32 | 4.35 |
| amine oxidase | AHJ39955.1 | 7 | 0.22 | 0.30 | 0.26 | 0.11 | 0.25 | 0.18 | 0.04 | 0.12 | 0.08 | 0.18 | 0.40 | 0.29 | 0.49 | 0.82 | 1.31 |
| hypothetical protein | AMK62059.1 | 33 | 7.98 | 10.41 | 9.20 | 4.58 | 6.76 | 5.67 | 1.52 | 3.62 | 2.57 | 0.19 | 0.35 | 0.27 | 0.57 | 0.65 | 1.22 |
| choline dehydrogenase-like protein | AMK62096.1 | 27 | 1.64 | 2.28 | 1.96 | 0.63 | 1.08 | 0.86 | 0.23 | 0.74 | 0.49 | 0.14 | 0.33 | 0.23 | 0.38 | 0.47 | 0.86 |
| hypothetical protein | AHJ40128.1 | 94 | 1.76 | 0.79 | 1.27 | 0.06 | 1.98 | 1.02 | 0.06 | 0.33 | 0.19 | 0.03 | 0.41 | 0.22 | 0.04 | 2.50 | 2.54 |
| hypothetical protein | AMK61902.1 | 9 | 0.56 | 0.65 | 0.61 | 0.67 | 0.55 | 0.61 | 0.00 | 0.29 | 0.15 | 0.00 | 0.45 | 0.22 | 1.20 | 0.84 | 2.04 |
| choline dehydrogenase-like protein | AMK61776.1 | 56 | 25.29 | 33.00 | 29.15 | 9.08 | 19.55 | 14.31 | 3.67 | 8.51 | 6.09 | 0.15 | 0.26 | 0.20 | 0.36 | 0.59 | 0.95 |
| hypothetical protein | AHJ40316.2 | 2 | 0.29 | 0.25 | 0.27 | 0.00 | 0.09 | 0.05 | 0.00 | 0.09 | 0.04 | 0.00 | 0.34 | 0.17 | 0.00 | 0.37 | 0.37 |
| hypothetical protein | AHJ40339.1 | 15 | 0.10 | 0.13 | 0.12 | 0.04 | 0.05 | 0.05 | 0.00 | 0.04 | 0.02 | 0.00 | 0.29 | 0.15 | 0.43 | 0.41 | 0.84 |
| WD repeat-containing protein | AHJ40002.1 | 17 | 0.33 | 0.27 | 0.30 | 0.05 | 0.25 | 0.15 | 0.00 | 0.07 | 0.04 | 0.00 | 0.27 | 0.14 | 0.17 | 0.95 | 1.11 |
| hypothetical protein | AHJ40318.2 | 10 | 0.08 | 0.15 | 0.11 | 0.03 | 0.36 | 0.19 | 0.00 | 0.04 | 0.02 | 0.00 | 0.25 | 0.13 | 0.31 | 2.41 | 2.72 |
| hypothetical protein | AMK61942.1 | 10 | 1.20 | 0.52 | 0.86 | 0.12 | 0.47 | 0.30 | 0.06 | 0.09 | 0.07 | 0.05 | 0.18 | 0.11 | 0.10 | 0.91 | 1.02 |
| hypothetical protein | AMK61856.1 | 23 | 0.88 | 0.99 | 0.94 | 0.63 | 0.63 | 0.63 | 0.00 | 0.16 | 0.08 | 0.00 | 0.16 | 0.08 | 0.71 | 0.64 | 1.35 |
| capsid protein 1 | AHJ40114.2 | 45 | 1.50 | 0.98 | 1.24 | 0.09 | 1.56 | 0.82 | 0.00 | 0.11 | 0.06 | 0.00 | 0.11 | 0.06 | 0.06 | 1.59 | 1.65 |
| collagen triple helix repeat containing protein | AMK61745.1 | 9 | 1.82 | 1.56 | 1.69 | 0.07 | 0.32 | 0.20 | 0.00 | 0.15 | 0.08 | 0.00 | 0.10 | 0.05 | 0.04 | 0.21 | 0.24 |
| GMC-type oxidoreductase | AMK61775.1 | 3 | 0.05 | 0.05 | 0.05 | 0.08 | 0.08 | 0.08 | 0.00 | 0.00 | 0.00 | 0.00 | 0.00 | 0.00 | 1.56 | 1.44 | 3.00 |
| glucose-methanol-choline oxidoreductase | AHJ40412.1 | 4 | 0.06 | 0.08 | 0.07 | 0.00 | 0.03 | 0.01 | 0.00 | 0.00 | 0.00 | 0.00 | 0.00 | 0.00 | 0.00 | 0.31 | 0.31 |
| collagen triple helix repeat containing protein | AHJ40289.2 | 2 | 0.09 | 0.14 | 0.11 | 0.00 | 0.10 | 0.05 | 0.00 | 0.00 | 0.00 | 0.00 | 0.00 | 0.00 | 0.00 | 0.71 | 0.71 |
| hypothetical protein | AHJ40232.2 | 3 | 0.04 | 0.04 | 0.04 | 0.00 | 0.03 | 0.02 | 0.00 | 0.00 | 0.00 | 0.00 | 0.00 | 0.00 | 0.00 | 0.78 | 0.78 |
| translocase of outer mitochondrial membrane 40 | ADZ24223.1 | 2 | 0.02 | 0.02 | 0.02 | 0.00 | 0.02 | 0.01 | 0.00 | 0.00 | 0.00 | 0.00 | 0.00 | 0.00 | 0.00 | 1.18 | 1.18 |
| putative lipoxygenase | AMK61740.1 | 5 | 0.08 | 0.08 | 0.08 | 0.00 | 0.06 | 0.03 | 0.00 | 0.00 | 0.00 | 0.00 | 0.00 | 0.00 | 0.00 | 0.67 | 0.67 |
| hypothetical protein | AMK61967.1 | 3 | 0.02 | 0.01 | 0.02 | 0.00 | 0.05 | 0.03 | 0.00 | 0.00 | 0.00 | 0.00 | 0.00 | 0.00 | 0.00 | 3.45 | 3.45 |
| hypothetical protein | AMK61977.1 | 4 | 0.04 | 0.07 | 0.06 | 0.11 | 0.16 | 0.14 | 0.00 | 0.00 | 0.00 | 0.00 | 0.00 | 0.00 | 2.94 | 2.22 | 5.15 |
| hypothetical protein | AMK61837.1 | 6 | 0.05 | 0.07 | 0.06 | 0.00 | 0.03 | 0.01 | 0.00 | 0.00 | 0.00 | 0.00 | 0.00 | 0.00 | 0.00 | 0.38 | 0.38 |
| hypothetical protein | AHJ40107.2 | 3 | 0.09 | 0.04 | 0.06 | 0.00 | 0.09 | 0.04 | 0.00 | 0.00 | 0.00 | 0.00 | 0.00 | 0.00 | 0.00 | 2.44 | 2.44 |
| DNA-dir. RNAP subunit RPB9 | AHJ39967.2 | 7 | 0.03 | 0.07 | 0.05 | 0.24 | 0.07 | 0.16 | 0.00 | 0.00 | 0.00 | 0.00 | 0.00 | 0.00 | 6.94 | 1.07 | 8.01 |
| mRNA-capping enzyme | AHJ40083.1 | 10 | 0.05 | 0.02 | 0.03 | 0.00 | 0.01 | 0.01 | 0.00 | 0.00 | 0.00 | 0.00 | 0.00 | 0.00 | 0.00 | 0.91 | 0.91 |
| DNA-dir. RNAP subunit 1 | AHJ40172.1 | 16 | 0.04 | 0.04 | 0.04 | 0.00 | 0.06 | 0.03 | 0.00 | 0.00 | 0.00 | 0.00 | 0.00 | 0.00 | 0.00 | 1.54 | 1.54 |
| prolyl 4-hydroxylase | AMK61959.1 | 6 | 0.05 | 0.05 | 0.05 | 0.00 | 0.06 | 0.03 | 0.00 | 0.00 | 0.00 | 0.00 | 0.00 | 0.00 | 0.00 | 1.10 | 1.10 |
| hypothetical protein | AMK61849.1 | 6 | 0.05 | 0.03 | 0.04 | 0.00 | 0.07 | 0.03 | 0.00 | 0.00 | 0.00 | 0.00 | 0.00 | 0.00 | 0.00 | 2.45 | 2.45 |
| hypothetical protein | AHJ40243.1 | 5 | 0.06 | 0.08 | 0.07 | 0.00 | 0.09 | 0.04 | 0.00 | 0.00 | 0.00 | 0.00 | 0.00 | 0.00 | 0.00 | 1.05 | 1.05 |
| hypothetical protein | AHJ40051.1 | 8 | 0.17 | 0.11 | 0.14 | 0.06 | 0.21 | 0.14 | 0.00 | 0.00 | 0.00 | 0.00 | 0.00 | 0.00 | 0.36 | 1.98 | 2.34 |
| hypothetical protein | AMK61892.1 | 17 | 0.08 | 0.05 | 0.06 | 0.00 | 0.15 | 0.08 | 0.00 | 0.00 | 0.00 | 0.00 | 0.00 | 0.00 | 0.00 | 3.18 | 3.18 |
| NHL repeat-containing protein | AMK61987.1 | 7 | 0.02 | 0.04 | 0.03 | 0.00 | 0.10 | 0.05 | 0.00 | 0.00 | 0.00 | 0.00 | 0.00 | 0.00 | 0.00 | 2.14 | 2.14 |
| alpha beta hydrolase/esterase/lipase | AHJ40190.1 | 10 | 0.13 | 0.12 | 0.13 | 0.00 | 0.12 | 0.06 | 0.00 | 0.00 | 0.00 | 0.00 | 0.00 | 0.00 | 0.00 | 1.05 | 1.05 |
| hypothetical protein | AHJ40144.1 | 9 | 0.07 | 0.04 | 0.05 | 0.35 | 0.16 | 0.25 | 0.00 | 0.00 | 0.00 | 0.00 | 0.00 | 0.00 | 5.32 | 4.44 | 9.76 |
| hypothetical protein | AHJ40220.1 | 18 | 0.24 | 0.09 | 0.16 | 0.30 | 0.24 | 0.27 | 0.00 | 0.00 | 0.00 | 0.00 | 0.00 | 0.00 | 1.28 | 2.69 | 3.97 |
| hypothetical protein | AHJ40061.1 | 22 | 0.03 | 0.02 | 0.02 | 0.00 | 0.03 | 0.01 | 0.00 | 0.00 | 0.00 | 0.00 | 0.00 | 0.00 | 0.00 | 1.78 | 1.78 |
| hypothetical protein | AHJ40254.1 | 5 | 0.05 | 0.06 | 0.06 | 0.06 | 0.11 | 0.08 | 0.00 | 0.00 | 0.00 | 0.00 | 0.00 | 0.00 | 1.13 | 1.70 | 2.82 |
| hypothetical protein | AHJ40060.1 | 4 | 0.06 | 0.07 | 0.07 | 0.00 | 0.06 | 0.03 | 0.00 | 0.00 | 0.00 | 0.00 | 0.00 | 0.00 | 0.00 | 0.88 | 0.88 |
| regulator of chromosome condensation | AMK61866.1 | 30 | 0.05 | 0.06 | 0.06 | 0.00 | 0.05 | 0.02 | 0.00 | 0.00 | 0.00 | 0.00 | 0.00 | 0.00 | 0.00 | 0.82 | 0.82 |
| ***Tupanvirus soda lake*** | | | | | | | | | | | | | | | | | |
| *Protein* | *Accession ID* | *Pep* | *Material Applied* | | | *Pellet* | | | *Supernatant* | | | *Supernatant/MA* | | | *Pellet/MA* | | |
|  |  |  | *2* | *3* | *Avg.* | *2* | *3* | *Avg.* | *2* | *3* | *Avg.* | *2* | *3* | *Avg.* | *2* | *3* | *Avg.* |
|  |  |  | ***%*** | ***%*** | ***%*** | ***%*** | ***%*** | ***%*** | ***%*** | ***%*** | ***%*** |  |  |  |  |  |  |
| actin | CAA23399.1 | 7 | 0.06 | 0.02 | 0.04 | 0.08 | 0.02 | 0.05 | 77.04 | 0.76 | 38.90 | 1308 | 42.18 | 675 | 1.28 | 0.93 | 1.10 |
| ubiquitin domain-containing protein | AUL78040.1 | 6 | 0.21 | 0.26 | 0.24 | 0.10 | 0.12 | 0.11 | 18.23 | 5.36 | 11.79 | 85.34 | 20.77 | 53.05 | 0.46 | 0.47 | 0.47 |
| putative ORFan | AUL78088.1 | 7 | 0.15 | 0.23 | 0.19 | 0.00 | 0.08 | 0.04 | 0.00 | 6.92 | 3.46 | 0.00 | 29.87 | 14.93 | 0.00 | 0.32 | 0.16 |
| hypothetical protein | AUL78468.1 | 3 | 0.07 | 0.09 | 0.08 | 0.03 | 0.04 | 0.03 | 0.00 | 1.85 | 0.93 | 0.00 | 20.37 | 10.19 | 0.40 | 0.46 | 0.43 |
| glutaredoxin | AUL78724.1 | 2 | 0.06 | 0.06 | 0.06 | 0.02 | 0.00 | 0.01 | 0.00 | 1.08 | 0.54 | 0.00 | 19.35 | 9.67 | 0.25 | 0.00 | 0.12 |
| hypothetical protein | AUL78348.1 | 3 | 0.01 | 0.01 | 0.01 | 0.00 | 0.00 | 0.00 | 0.00 | 0.09 | 0.05 | 0.00 | 13.75 | 6.87 | 0.00 | 0.00 | 0.00 |
| DNA-dir. RNAP. subunit | AUL78016.1 | 3 | 0.08 | 0.06 | 0.07 | 0.07 | 0.08 | 0.08 | 0.00 | 0.73 | 0.37 | 0.00 | 12.91 | 6.45 | 0.91 | 1.37 | 1.14 |
| hypothetical protein | AUL78055.1 | 5 | 0.12 | 0.13 | 0.12 | 0.06 | 0.06 | 0.06 | 0.00 | 1.52 | 0.76 | 0.00 | 11.56 | 5.78 | 0.47 | 0.43 | 0.45 |
| hypothetical protein | AUL77930.1 | 3 | 0.05 | 0.04 | 0.04 | 0.00 | 0.00 | 0.00 | 0.00 | 0.35 | 0.18 | 0.00 | 9.77 | 4.88 | 0.00 | 0.00 | 0.00 |
| hypothetical protein | AUL78681.1 | 2 | 0.13 | 0.21 | 0.17 | 0.03 | 0.03 | 0.03 | 0.00 | 2.01 | 1.01 | 0.00 | 9.55 | 4.78 | 0.19 | 0.15 | 0.17 |
| DNA--dir. RNAP subunit 6 | AUL78368.1 | 5 | 0.09 | 0.03 | 0.06 | 0.00 | 0.01 | 0.00 | 0.00 | 0.21 | 0.10 | 0.00 | 7.76 | 3.88 | 0.00 | 0.33 | 0.17 |
| hypothetical protein | AUL77723.1 | 8 | 0.40 | 0.42 | 0.41 | 0.30 | 0.26 | 0.28 | 0.00 | 1.99 | 1.00 | 0.00 | 4.78 | 2.39 | 0.75 | 0.62 | 0.69 |
| hypothetical protein | AUL77907.1 | 9 | 0.69 | 0.61 | 0.65 | 0.49 | 0.62 | 0.55 | 0.00 | 1.85 | 0.92 | 0.00 | 3.03 | 1.51 | 0.71 | 1.01 | 0.86 |
| hypothetical protein | AUL78288.1 | 5 | 0.74 | 0.52 | 0.63 | 0.31 | 0.17 | 0.24 | 0.00 | 1.44 | 0.72 | 0.00 | 2.74 | 1.37 | 0.41 | 0.32 | 0.37 |
| hypothetical protein | AUL78466.1 | 4 | 0.09 | 0.07 | 0.08 | 0.07 | 0.03 | 0.05 | 0.00 | 0.17 | 0.08 | 0.00 | 2.45 | 1.23 | 0.85 | 0.37 | 0.61 |
| mg709 protein | AUL77661.1 | 4 | 0.42 | 0.33 | 0.38 | 0.10 | 0.18 | 0.14 | 0.00 | 0.64 | 0.32 | 0.00 | 1.93 | 0.97 | 0.25 | 0.56 | 0.40 |
| hypothetical protein | AUL78191.1 | 31 | 8.73 | 9.74 | 9.23 | 5.43 | 8.84 | 7.13 | 0.00 | 18.59 | 9.29 | 0.00 | 1.91 | 0.95 | 0.62 | 0.91 | 0.76 |
| putative pore coat assembly factor | AUL78211.1 | 11 | 0.31 | 0.36 | 0.34 | 0.21 | 0.23 | 0.22 | 0.00 | 0.64 | 0.32 | 0.00 | 1.75 | 0.88 | 0.68 | 0.65 | 0.66 |
| catalase HPII | AUL78097.1 | 12 | 0.66 | 0.74 | 0.70 | 0.44 | 0.52 | 0.48 | 0.00 | 1.08 | 0.54 | 0.00 | 1.45 | 0.73 | 0.66 | 0.70 | 0.68 |
| thioredoxin domain-containing protein | AUL77963.1 | 9 | 1.75 | 1.75 | 1.75 | 0.61 | 0.34 | 0.48 | 0.00 | 2.38 | 1.19 | 0.00 | 1.36 | 0.68 | 0.35 | 0.20 | 0.27 |
| hypothetical protein | AUL77936.1 | 6 | 0.17 | 0.28 | 0.22 | 0.33 | 0.29 | 0.31 | 0.00 | 0.34 | 0.17 | 0.00 | 1.23 | 0.62 | 1.94 | 1.06 | 1.50 |
| putative protein kinase | AUL78629.1 | 4 | 0.54 | 0.33 | 0.44 | 0.40 | 0.22 | 0.31 | 0.00 | 0.22 | 0.11 | 0.00 | 0.66 | 0.33 | 0.74 | 0.67 | 0.71 |
| DNA-dir. RNAP subunit 1 | AUL78302.1 | 32 | 0.24 | 0.60 | 0.42 | 0.33 | 0.81 | 0.57 | 0.00 | 0.39 | 0.20 | 0.00 | 0.65 | 0.32 | 1.38 | 1.35 | 1.36 |
| arylsulfatase | AUL78269.1 | 9 | 0.39 | 0.42 | 0.40 | 0.47 | 0.33 | 0.40 | 0.00 | 0.27 | 0.14 | 0.00 | 0.64 | 0.32 | 1.22 | 0.78 | 1.00 |
| kinesin-like protein | AUL77838.1 | 12 | 0.15 | 0.13 | 0.14 | 0.20 | 0.05 | 0.12 | 0.00 | 0.08 | 0.04 | 0.00 | 0.61 | 0.31 | 1.34 | 0.35 | 0.85 |
| hypothetical protein | AUL77694.1 | 8 | 1.56 | 2.58 | 2.07 | 1.36 | 2.90 | 2.13 | 0.00 | 1.57 | 0.78 | 0.00 | 0.61 | 0.30 | 0.87 | 1.13 | 1.00 |
| capsid protein 1 | AUL78147.1 | 47 | 46.74 | 42.28 | 44.51 | 55.65 | 51.77 | 53.71 | 4.73 | 20.72 | 12.72 | 0.10 | 0.49 | 0.30 | 1.19 | 1.22 | 1.21 |
| hypothetical protein | AUL78067.1 | 10 | 0.32 | 0.32 | 0.32 | 0.39 | 0.31 | 0.35 | 0.00 | 0.16 | 0.08 | 0.00 | 0.48 | 0.24 | 1.23 | 0.96 | 1.09 |
| major core protein | AUL78082.1 | 35 | 5.43 | 6.32 | 5.88 | 7.44 | 4.84 | 6.14 | 0.00 | 2.92 | 1.46 | 0.00 | 0.46 | 0.23 | 1.37 | 0.77 | 1.07 |
| hypothetical protein | AUL78214.1 | 10 | 1.34 | 1.23 | 1.28 | 0.94 | 1.31 | 1.13 | 0.00 | 0.56 | 0.28 | 0.00 | 0.46 | 0.23 | 0.70 | 1.06 | 0.88 |
| putative fibril associated protein | AUL78400.1 | 17 | 5.65 | 4.00 | 4.83 | 5.05 | 4.99 | 5.02 | 0.00 | 1.72 | 0.86 | 0.00 | 0.43 | 0.22 | 0.89 | 1.25 | 1.07 |
| hypothetical protein | AUL78287.1 | 14 | 1.78 | 2.35 | 2.07 | 0.77 | 1.16 | 0.97 | 0.00 | 1.00 | 0.50 | 0.00 | 0.42 | 0.21 | 0.43 | 0.49 | 0.46 |
| hypothetical protein | AUL78232.1 | 3 | 0.28 | 0.19 | 0.23 | 0.03 | 0.08 | 0.05 | 0.00 | 0.07 | 0.03 | 0.00 | 0.38 | 0.19 | 0.11 | 0.40 | 0.26 |
| hypothetical protein | AUL78219.1 | 8 | 1.62 | 2.22 | 1.92 | 1.90 | 2.55 | 2.22 | 0.00 | 0.83 | 0.41 | 0.00 | 0.37 | 0.19 | 1.17 | 1.15 | 1.16 |
| putative ORFan | AUL77729.1 | 7 | 0.76 | 0.50 | 0.63 | 0.40 | 0.75 | 0.57 | 0.00 | 0.16 | 0.08 | 0.00 | 0.33 | 0.16 | 0.53 | 1.51 | 1.02 |
| hypothetical protein | AUL77688.1 | 5 | 3.52 | 3.90 | 3.71 | 2.37 | 3.13 | 2.75 | 0.00 | 1.18 | 0.59 | 0.00 | 0.30 | 0.15 | 0.67 | 0.80 | 0.74 |
| hypothetical protein | AUL78135.1 | 11 | 2.42 | 2.17 | 2.29 | 2.03 | 1.62 | 1.83 | 0.00 | 0.64 | 0.32 | 0.00 | 0.30 | 0.15 | 0.84 | 0.75 | 0.79 |
| hypothetical protein | AUL78143.1 | 9 | 5.33 | 7.20 | 6.26 | 5.91 | 4.91 | 5.41 | 0.00 | 2.07 | 1.04 | 0.00 | 0.29 | 0.14 | 1.11 | 0.68 | 0.90 |
| intein-containing DNA-dir. RNAP subunit 2 | AUL78362.1 | 7 | 0.17 | 0.24 | 0.20 | 0.18 | 0.26 | 0.22 | 0.00 | 0.06 | 0.03 | 0.00 | 0.24 | 0.12 | 1.08 | 1.11 | 1.09 |
| hypothetical protein | AUL77600.1 | 3 | 0.39 | 0.34 | 0.37 | 0.46 | 0.20 | 0.33 | 0.00 | 0.06 | 0.03 | 0.00 | 0.16 | 0.08 | 1.18 | 0.59 | 0.88 |
| hypothetical protein | AUL78481.1 | 4 | 0.07 | 0.17 | 0.12 | 0.08 | 0.14 | 0.11 | 0.00 | 0.02 | 0.01 | 0.00 | 0.14 | 0.07 | 1.15 | 0.84 | 0.99 |
| hypothetical protein | AUL77492.1 | 3 | 0.01 | 0.02 | 0.02 | 0.01 | 0.02 | 0.01 | 0.00 | 0.00 | 0.00 | 0.00 | 0.00 | 0.00 | 1.12 | 0.72 | 0.92 |
| hypothetical protein | AUL77863.1 | 2 | 0.02 | 0.03 | 0.03 | 0.01 | 0.03 | 0.02 | 0.00 | 0.00 | 0.00 | 0.00 | 0.00 | 0.00 | 0.61 | 0.82 | 0.71 |
| DNA-dir. RNAP subunit | AUL78244.1 | 3 | 0.02 | 0.03 | 0.02 | 0.07 | 0.03 | 0.05 | 0.00 | 0.00 | 0.00 | 0.00 | 0.00 | 0.00 | 3.25 | 0.97 | 2.11 |
| thiol oxidoreductase E10R | AUL77655.1 | 2 | 0.02 | 0.01 | 0.02 | 0.03 | 0.01 | 0.02 | 0.00 | 0.00 | 0.00 | 0.00 | 0.00 | 0.00 | 1.32 | 0.79 | 1.06 |
| putative ankyrin repeat protein | AUL78278.1 | 6 | 0.02 | 0.02 | 0.02 | 0.05 | 0.03 | 0.04 | 0.00 | 0.00 | 0.00 | 0.00 | 0.00 | 0.00 | 1.87 | 1.29 | 1.58 |
| bifunctional metalloprotease ubiquitin-protein ligase | AUL78691.1 | 3 | 0.03 | 0.05 | 0.04 | 0.04 | 0.05 | 0.04 | 0.00 | 0.00 | 0.00 | 0.00 | 0.00 | 0.00 | 1.17 | 0.91 | 1.04 |
| hypothetical protein | AUL78731.1 | 2 | 0.03 | 0.02 | 0.02 | 0.00 | 0.00 | 0.00 | 0.00 | 0.00 | 0.00 | 0.00 | 0.00 | 0.00 | 0.00 | 0.00 | 0.00 |
| putative ORFan | AUL77532.1 | 2 | 0.03 | 0.03 | 0.03 | 0.05 | 0.03 | 0.04 | 0.00 | 0.00 | 0.00 | 0.00 | 0.00 | 0.00 | 1.75 | 0.87 | 1.31 |
| hypothetical protein | AUL78045.1 | 3 | 0.03 | 0.08 | 0.05 | 0.03 | 0.08 | 0.06 | 0.00 | 0.00 | 0.00 | 0.00 | 0.00 | 0.00 | 1.03 | 1.00 | 1.01 |
| structural ppiase-like protein | AUL77649.1 | 4 | 0.04 | 0.03 | 0.03 | 0.02 | 0.03 | 0.03 | 0.00 | 0.00 | 0.00 | 0.00 | 0.00 | 0.00 | 0.68 | 0.84 | 0.76 |
| hypothetical protein | AUL77666.1 | 5 | 0.04 | 0.03 | 0.03 | 0.12 | 0.03 | 0.07 | 0.00 | 0.00 | 0.00 | 0.00 | 0.00 | 0.00 | 3.01 | 1.11 | 2.06 |
| mg749 protein | AUL77517.1 | 1 | 0.04 | 0.04 | 0.04 | 0.02 | 0.02 | 0.02 | 0.00 | 0.00 | 0.00 | 0.00 | 0.00 | 0.00 | 0.42 | 0.47 | 0.44 |
| hypothetical protein | AUL78068.1 | 7 | 0.04 | 0.03 | 0.04 | 0.05 | 0.04 | 0.05 | 0.00 | 0.00 | 0.00 | 0.00 | 0.00 | 0.00 | 1.12 | 1.37 | 1.25 |
| dna topoisomerase 1b | AUL78109.1 | 10 | 0.05 | 0.06 | 0.05 | 0.03 | 0.05 | 0.04 | 0.00 | 0.00 | 0.00 | 0.00 | 0.00 | 0.00 | 0.72 | 0.91 | 0.81 |
| intein-containing DNA-dir. RNAP subunit 2 | AUL78361.1 | 12 | 0.05 | 0.09 | 0.07 | 0.11 | 0.11 | 0.11 | 0.00 | 0.00 | 0.00 | 0.00 | 0.00 | 0.00 | 2.23 | 1.29 | 1.76 |
| chemotaxis | AUL78637.1 | 4 | 0.05 | 0.06 | 0.06 | 0.06 | 0.06 | 0.06 | 0.00 | 0.00 | 0.00 | 0.00 | 0.00 | 0.00 | 1.25 | 0.95 | 1.10 |
| phosphoesterase-like protein | AUL77796.1 | 4 | 0.05 | 0.12 | 0.09 | 0.05 | 0.08 | 0.07 | 0.00 | 0.00 | 0.00 | 0.00 | 0.00 | 0.00 | 0.95 | 0.67 | 0.81 |
| hypothetical protein | AUL78280.1 | 2 | 0.05 | 0.03 | 0.04 | 0.05 | 0.04 | 0.04 | 0.00 | 0.00 | 0.00 | 0.00 | 0.00 | 0.00 | 0.96 | 1.39 | 1.17 |
| hypothetical protein | AUL78198.1 | 2 | 0.06 | 0.05 | 0.05 | 0.00 | 0.03 | 0.01 | 0.00 | 0.00 | 0.00 | 0.00 | 0.00 | 0.00 | 0.00 | 0.57 | 0.28 |
| hypothetical protein | AUL77647.1 | 7 | 0.06 | 0.04 | 0.05 | 0.05 | 0.05 | 0.05 | 0.00 | 0.00 | 0.00 | 0.00 | 0.00 | 0.00 | 0.92 | 1.06 | 0.99 |
| hypothetical protein | AUL78155.1 | 2 | 0.06 | 0.13 | 0.09 | 0.10 | 0.08 | 0.09 | 0.00 | 0.00 | 0.00 | 0.00 | 0.00 | 0.00 | 1.59 | 0.66 | 1.12 |
| hypothetical protein | AUL78061.1 | 3 | 0.07 | 0.01 | 0.04 | 0.00 | 0.00 | 0.00 | 0.00 | 0.00 | 0.00 | 0.00 | 0.00 | 0.00 | 0.00 | 0.00 | 0.00 |
| putative protein phosphatase 2c | AUL77859.1 | 4 | 0.07 | 0.08 | 0.07 | 0.07 | 0.08 | 0.08 | 0.00 | 0.00 | 0.00 | 0.00 | 0.00 | 0.00 | 1.03 | 1.06 | 1.05 |
| FtsJ-like methyl transferase | AUL78032.1 | 5 | 0.07 | 0.07 | 0.07 | 0.09 | 0.09 | 0.09 | 0.00 | 0.00 | 0.00 | 0.00 | 0.00 | 0.00 | 1.40 | 1.42 | 1.41 |
| hypothetical protein | AUL77903.1 | 3 | 0.08 | 0.06 | 0.07 | 0.09 | 0.06 | 0.07 | 0.00 | 0.00 | 0.00 | 0.00 | 0.00 | 0.00 | 1.05 | 1.02 | 1.03 |
| hypothetical protein | AUL77961.1 | 5 | 0.08 | 0.08 | 0.08 | 0.08 | 0.06 | 0.07 | 0.00 | 0.00 | 0.00 | 0.00 | 0.00 | 0.00 | 0.98 | 0.79 | 0.88 |
| SNF2 family helicase | AUL77941.1 | 9 | 0.08 | 0.06 | 0.07 | 0.15 | 0.15 | 0.15 | 0.00 | 0.00 | 0.00 | 0.00 | 0.00 | 0.00 | 1.77 | 2.43 | 2.10 |
| polyA polymerase catalitic subunit | AUL77929.1 | 12 | 0.08 | 0.18 | 0.13 | 0.09 | 0.19 | 0.14 | 0.00 | 0.00 | 0.00 | 0.00 | 0.00 | 0.00 | 1.12 | 1.03 | 1.08 |
| hypothetical protein | AUL78093.1 | 4 | 0.10 | 0.23 | 0.16 | 0.07 | 0.16 | 0.12 | 0.00 | 0.00 | 0.00 | 0.00 | 0.00 | 0.00 | 0.74 | 0.71 | 0.73 |
| hypothetical protein | AUL78319.1 | 4 | 0.10 | 0.07 | 0.09 | 0.10 | 0.07 | 0.09 | 0.00 | 0.00 | 0.00 | 0.00 | 0.00 | 0.00 | 1.02 | 0.97 | 1.00 |
| thioredoxin domain-containing protein | AUL78192.1 | 2 | 0.11 | 0.11 | 0.11 | 0.17 | 0.00 | 0.08 | 0.00 | 0.00 | 0.00 | 0.00 | 0.00 | 0.00 | 1.50 | 0.00 | 0.75 |
| DNA-dep. RNAP subunit Rpb9 | AUL78739.1 | 4 | 0.12 | 0.10 | 0.11 | 0.12 | 0.07 | 0.09 | 0.00 | 0.00 | 0.00 | 0.00 | 0.00 | 0.00 | 0.99 | 0.67 | 0.83 |
| hypothetical protein | AUL77933.1 | 6 | 0.12 | 0.15 | 0.14 | 0.14 | 0.14 | 0.14 | 0.00 | 0.00 | 0.00 | 0.00 | 0.00 | 0.00 | 1.17 | 0.90 | 1.04 |
| mRNA capping enzyme | AUL78031.1 | 9 | 0.13 | 0.17 | 0.15 | 0.16 | 0.22 | 0.19 | 0.00 | 0.00 | 0.00 | 0.00 | 0.00 | 0.00 | 1.19 | 1.33 | 1.26 |
| glycosyl hydrolase family 18 | AUL77711.1 | 2 | 0.13 | 0.13 | 0.13 | 0.09 | 0.13 | 0.11 | 0.00 | 0.00 | 0.00 | 0.00 | 0.00 | 0.00 | 0.68 | 0.99 | 0.84 |
| NTPase | AUL78021.1 | 14 | 0.19 | 0.16 | 0.18 | 0.18 | 0.18 | 0.18 | 0.00 | 0.00 | 0.00 | 0.00 | 0.00 | 0.00 | 0.92 | 1.12 | 1.02 |
| putative oxireductase | AUL77599.1 | 8 | 0.20 | 0.29 | 0.24 | 0.14 | 0.18 | 0.16 | 0.00 | 0.00 | 0.00 | 0.00 | 0.00 | 0.00 | 0.70 | 0.63 | 0.66 |
| hypothetical protein | AUL78246.1 | 15 | 0.21 | 0.21 | 0.21 | 0.34 | 0.29 | 0.32 | 0.00 | 0.00 | 0.00 | 0.00 | 0.00 | 0.00 | 1.59 | 1.36 | 1.48 |
| putative ORFan | AUL78635.1 | 3 | 0.25 | 0.24 | 0.25 | 0.08 | 0.23 | 0.15 | 0.00 | 0.00 | 0.00 | 0.00 | 0.00 | 0.00 | 0.30 | 0.96 | 0.63 |
| hypothetical protein | AUL78601.1 | 4 | 0.27 | 0.09 | 0.18 | 0.14 | 0.09 | 0.11 | 0.00 | 0.00 | 0.00 | 0.00 | 0.00 | 0.00 | 0.51 | 1.03 | 0.77 |
| putative early transcription factor | AUL77899.1 | 19 | 0.28 | 0.42 | 0.35 | 0.35 | 0.38 | 0.36 | 0.00 | 0.00 | 0.00 | 0.00 | 0.00 | 0.00 | 1.22 | 0.90 | 1.06 |
| putative ORFan | AUL78206.1 | 4 | 0.39 | 0.49 | 0.44 | 0.00 | 0.00 | 0.00 | 0.00 | 0.00 | 0.00 | 0.00 | 0.00 | 0.00 | 0.00 | 0.00 | 0.00 |
| hypothetical protein | AUL77752.1 | 9 | 0.54 | 0.41 | 0.48 | 0.34 | 0.67 | 0.50 | 0.00 | 0.00 | 0.00 | 0.00 | 0.00 | 0.00 | 0.63 | 1.62 | 1.13 |
| hypothetical protein | AUL78292.1 | 3 | 0.67 | 0.49 | 0.58 | 0.07 | 0.17 | 0.12 | 0.00 | 0.00 | 0.00 | 0.00 | 0.00 | 0.00 | 0.11 | 0.35 | 0.23 |
