## Supplementary figures and images for "Boiling Acid Mimics Intracellular Giant Virus Genome Release"

### Supplemental Figure 1

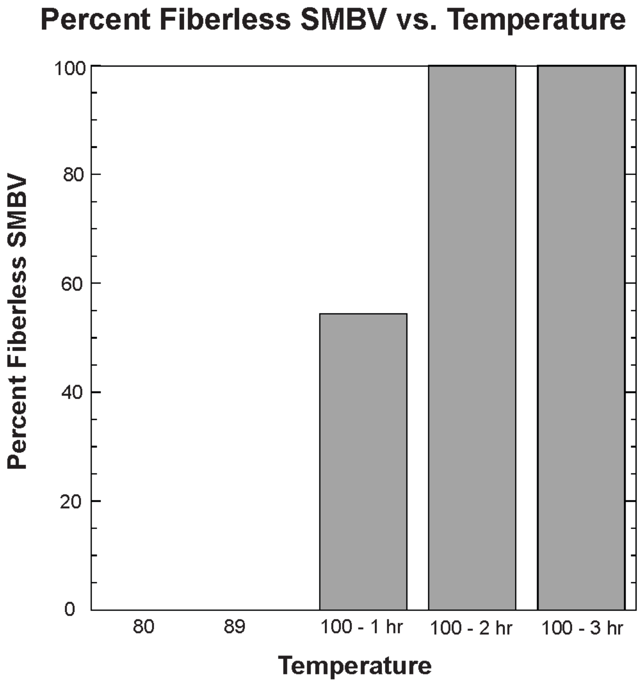

### Supplemental Figure 2

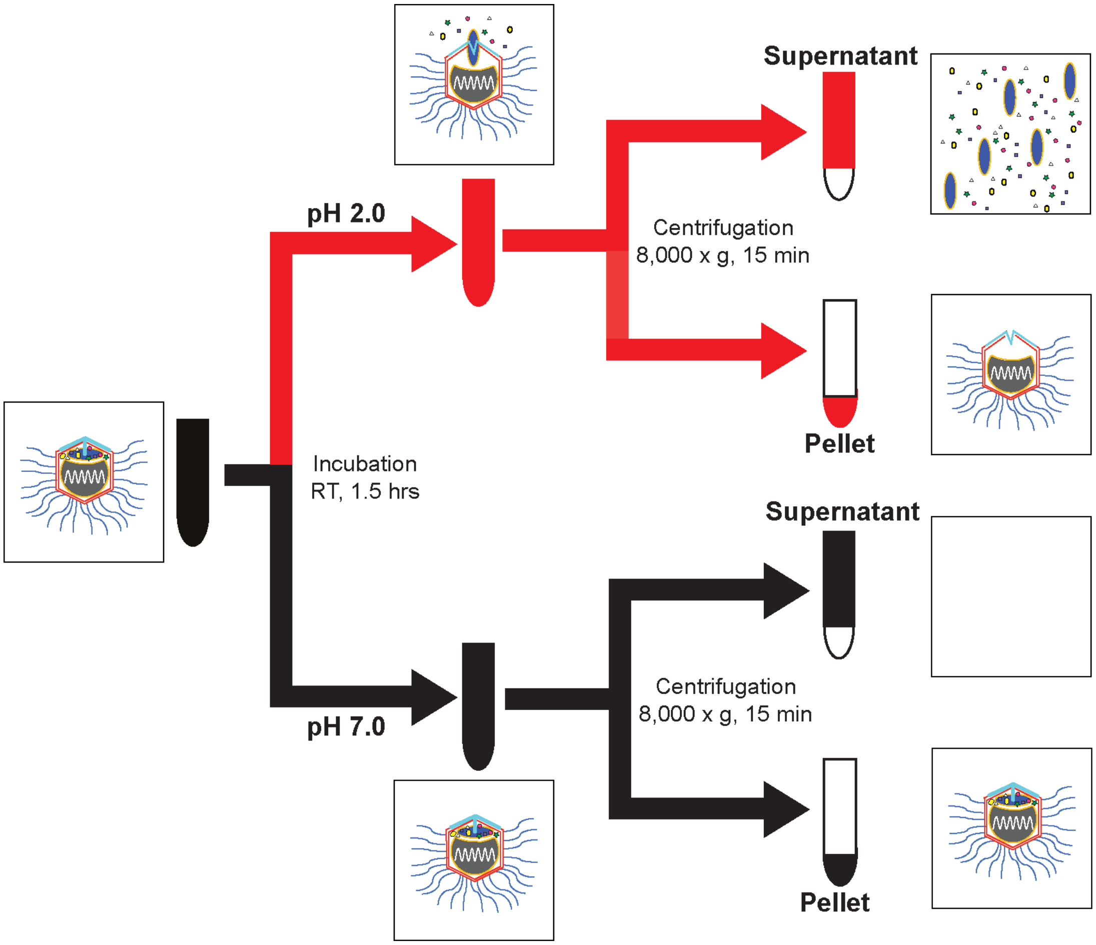
